# Liriodendrin Targets PFKFB3 to Suppress Inflammatory Phenotypic Transition and Vascular Remodeling in Pulmonary Hypertension

**DOI:** 10.64898/2026.07.30.741507

**Authors:** Qingye Zeng, Zhenzhen Duan, Qian Liu, Lei Yang, Zhao Sha, Yicheng Lv, Xiaoxun Huang, Jincheng Zhang, Jingyang Su, Zhiqian Lu, Shaowen Liu, Deping Kong

**Affiliations:** Department of Cardiology, Shanghai General Hospital, School of Medicine, Shanghai Jiao Tong University, Shanghai, China; Precision Research Center for Refractory Diseases, Institute for Clinical Research, Shanghai General Hospital, School of Medicine, Shanghai Jiao Tong University, Shanghai, China; Tianjin NanKai Hospital, Tianjin Medical University, Tianjin, 300100, China; Department of Clinical Pharmacology, School of Pharmacy, China Medical University, Shenyang Liaoning, 110122, China; Department of Cardiovascular Surgery, Shanghai Sixth People’s Hospital Affiliated to Shanghai Jiao Tong University School of Medicine, Shanghai, China.

**Keywords:** pulmonary hypertension, liriodendrin, PFKFB3, protein degradation, histone lactylation

## Abstract

**Background:** Pulmonary hypertension (PH) involves progressive vascular remodeling and perivascular inflammation. Despite modest clinical improvements with current therapies, their limited ability to reverse remodeling or restore immune homeostasis highlights the need for novel agents. *Liriodendrin* (Lidd), derived from *Sargentodoxae caulis*, exhibits anti-inflammatory and antiproliferative activities, but its efficacy and molecular targets in PH are unknown.

**Methods:** Two well-established PH animal models - the SU5416/hypoxia (SuHx) mice model and monocrotaline (MCT)-induced rat model - were employed for *in vivo* assessment of Lidd conducted pharmacological effects. Primary human pulmonary artery smooth muscle cells (hPASMCs) were utilized for mechanistic investigations. RNA-sequencing (RNA-seq) analysis was conducted to identify potential signaling pathways modulated by Lidd treatment. The direct molecular target of Lidd was determined through integrated application of drug affinity responsive target stability (DARTS) assay coupled with western blot validation. To delineate histone lactylation-mediated transcriptional regulation, we combined Cleavage Under Targets and Tagmentation (CUT&Tag) sequencing data analysis followed by chromatin immunoprecipitation quantitative PCR (ChIP-qPCR) verification. Genetic validation was achieved using PFKFB3-deficient murine models to verify the specificity of Lidd-mediated pharmacological actions.

**Results:** Lidd administration attenuated pulmonary vascular remodeling, perivascular macrophage infiltration and PH progression in both SuHx and MCT models. Transcriptomic profiling of Lidd-treated hPASMCs revealed predominant enrichment of downregulated genes in inflammatory and cytokine-associated pathways. Mechanistically, Lidd directly bound PFKFB3 and enhanced its interaction with FZR1, promoting PFKFB3 ubiquitination and degradation, which reduced glycolysis-driven lactate and consequent histone lactylation. This, in turn, diminished transcriptional activation of proliferative and inflammatory mediators, including CCND1, TNC, and CCL2. Notably, exogenous lactate supplementation or endogenous lactate accumulation restored histone lactylation and paradoxically potentiated Lidd’s inhibitory effects on PASMC proliferation and migration, whereas p300 inhibition abrogated these lactate-mediated effects. Importantly, Lidd failed to confer additional protection in PFKFB3-deficient mice, confirming PFKFB3 as the primary target mediating its therapeutic action.

**Conclusion:** Our findings reveal that Lidd selectively targets the PFKFB3-mediated glycolytic-epigenetic axis to suppress PASMC phenotypic transformation and pulmonary vascular remodeling, positioning it as a promising therapeutic candidate for PH.

## Introduction

Pulmonary hypertension (PH) represents a life-threatening cardiopulmonary disorder pathologically characterized by progressive pulmonary arterial remodeling, which triggers a cascade of hemodynamic consequences, culminating in elevated pulmonary vascular resistance and ultimately progressing to right ventricular failure^1,2^. Current treatment strategies, including endothelin receptor antagonists, phosphodiesterase inhibitors, and prostacyclin analogs, primarily target pathological vasoconstriction, yet exhibit limited efficacy in reversing established vascular remodeling^3,4^. Emerging evidence have revealed inflammation and autoimmune dysregulation as central drivers of vascular remodeling^5^. Clinical-pathological correlations demonstrate elevated circulating levels of pro-inflammatory mediators, including IL-6, TNF-α, and CCL2, were not only correlated with hemodynamic severity indices^5–10^, but also directly mediated endothelial dysfunction and phenotypic switching of pulmonary artery smooth muscle cells (PASMCs) through paracrine signaling^11–13^. Noteworthy, while immunomodulatory therapies show translational promise in preclinical models, critical individualized efficacy variations, long-term safety profiles, and precision targeting of specific immune pathways require systematic validation through large-scale randomized controlled trials^14–16^.

Beyond the well-documented pathological features of aberrant immune cell migration and infiltration, imbalance between pro-inflammatory and anti-inflammatory homeostasis, emerging research highlights the pivotal role of dysregulated stromal-immune cell interactions in shaping the vascular remodeling microenvironment^17–19^. Intriguingly, single-cell analyses reveal that normal pulmonary artery vessel walls and surrounding regions harbor innate immune cell populations, which maintain homeostatic regulation with structural cells via ligand-receptor networks^20^. However, signaling cascades become dominated by structural cell-derived mediators in PH. Mounting evidence demonstrates that vascular endothelial cells, smooth muscle cells, and fibroblasts acquire the capability to secrete core inflammatory mediators (e.g., IL-6/MCP-1), forming autocrine-paracrine positive feedback loops that ultimately drive irreversible pulmonary vascular remodeling^11,21^. Significantly, pulmonary arterial smooth muscle cells (PASMCs), the central components of the vascular medial layer, have uncovered not only exhibit characteristic contractile-synthetic phenotypic switching but also display striking functional heterogeneity^20,22,23^, including inflammatory phenotypic transition. Recent transcriptomic profiling reveals their concurrent upregulation of multiple inflammatory cytokine receptors (e.g., IL-8R) and ligands (e.g., CXCL12) in PASMCs, indicating their transformation from passive responders to active orchestrators of vascular remodeling^20^. But the mechanisms underlying PASMCs-immune cell communication networks and therapeutic targets are still not completely understood.

*Sargentodoxa cuneata (S. cuneata)*, a traditional medicinal plant, shows therapeutic potential in treating inflammatory and vascular diseases through its key bioactive constituent, Liriodendrin (Lidd)^24^. Lidd exerts anti-inflammatory, anti-proliferative, and vasoprotective effects by inhibiting inflammatory cytokine release, promoting nitric oxide (NO) via arginine metabolism, and enhancing cGMP signaling through phosphodiesterase (PDE) inhibition^25–28^. Lidd also regulates vascular permeability and endothelial function via the MYLK-Piezo1 pathway^29^, suggesting therapeutic potential for PH, but its precise role remains unclear.

In this study, we identified Lidd as a PH therapeutic candidate that targets PFKFB3-mediated glycolysis in PASMCs. Lidd significantly inhibited PDGF-BB-induced PASMCs proliferation, migration and pro-inflammatory cytokine secretion, thereby mitigating vascular remodeling. Mechanistically, Lidd interacted with PFKFB3 to promote its ubiquitination and degradation, reducing glycolytic flux, lactate production, and histone lactylation - key drivers of PH progression. Our findings highlight Lidd’s capacity to regulate PFKFB3 and glycolysis, inhibits PASMCs phenotypic transformation and inflammation, and ultimately modulates vascular remodeling, underscoring its potential as a novel PH therapy.

## Results

### Lidd prevents the PH Progression and Vascular Remodeling in Mice PH Model

Lidd is one of the four principal compounds (**Table S1**) identified in *Sargentodoxae Caulis*^30^. Lidd exhibited potential anti-hyperproliferative activity in addition to vasorelaxant effects, as indicated by systematic bioactivity predictions (**Table S2**). Specifically, Lidd was predicted to act as a cyclic AMP phosphodiesterase inhibitor and an antineoplastic agent by Prediction of Activity Spectra for Substances (PASS) (**Table S2**). Notably, in PDGF-BB-induced (20 ug/ml) proliferation models, Lidd administration (100 μM) revealed significant growth inhibition, whereas other components showed negligible effects (**Figure S1A-1B**). The same effect was observed in hypoxia-induced PASMCs proliferation (**Figure S1C-S1D**), suggesting its promise as a therapeutic candidate for vascular remodeling.

To explore the role of Lidd in the development of PH, we examined pulmonary hemodynamic and histological changes in SU5416/hypoxia (SuHx)–exposed mice after Lidd treatment (**Figure 1A**). Lidd dramatically attenuated the PH by reducing right ventricular systolic pressure (RVSP) (Lidd, 21.55±1.036 mm Hg versus vehicle, 28.00 ±1.204 mm Hg; P=0.0016; **Figure 1B-1C**) and ratio of the weight of the right ventricle to the weight of the left ventricle plus septum (RV/LV+S) (Lidd, 0.258±0.022 versus vehicle, 0.358±0.022; P=0.0013; **Figure 1D**). Furthermore, Lidd treatment led to an alleviated pathological vascular remodeling, as indicated by decreased pulmonary vascular wall thickness (media/cross-sectional area; **Figure 1E, G-H**), α-SMA (α-smooth muscle actin) expression (**Figure 1E**) and muscularization (**Figure 1F**). Interestingly, Lidd treatment markedly inhibited SuHx-induced PASMC proliferation by PCNA and α-SMA double staining (**Figure 1I-1J**). Additionally, Lidd treatment also protected against PH-induced right ventricular (RV) remodeling by decreasing cardiomyocyte cross-sectional area and collagen deposition in the RV (**Figure S2**). These results indicate that Lidd administration prevents SuHx-induced PH progression in mice.

**Figure 1.**
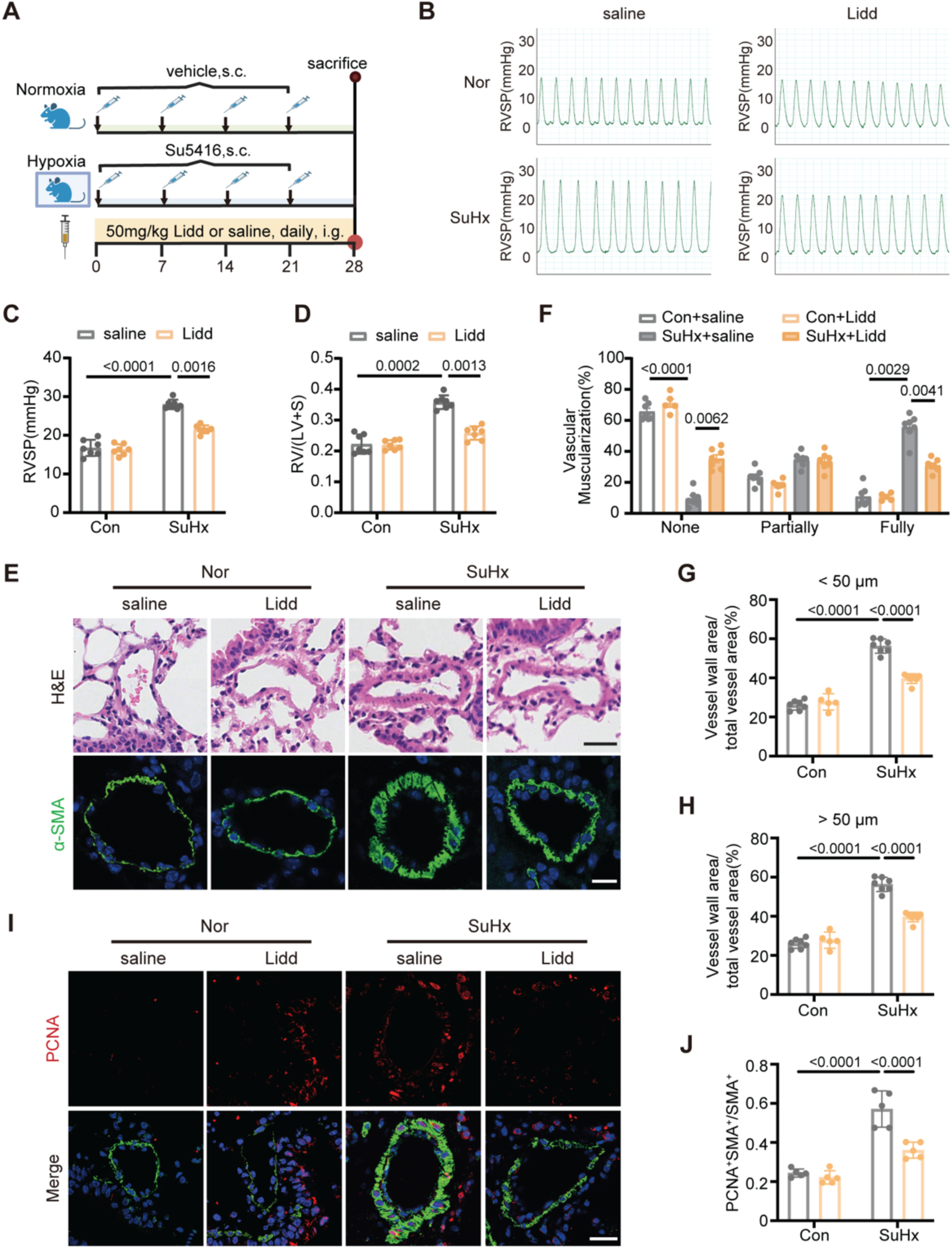
Liriodendrin (Lidd) attenuates SU5416/hypoxia (SuHx)-induced pulmonary hypertension (PH) in mice. **A**) Experimental design of the SuHx-induced PH mouse model. Mice were exposed to normoxia or SuHx for 28 days and received daily oral gavage of Lidd (50 mg/kg body weight) or saline throughout the experimental period. Hemodynamic measurements and tissue collection were performed on day 28. **B-C**) Representative right ventricular systolic pressure (RVSP) tracings (**B**) and quantification of RVSP (**C**) in control and SuHx mice treated with saline or Lidd (n = 7). **D)** Quantification of right ventricular hypertrophy, assessed by the ratio of right ventricular weight to left ventricular plus septal weight (RV/LV+S) (n = 7). **E)** Representative images of hematoxylin and eosin (H&E) staining (**top**) and α-smooth muscle actin (α-SMA, green) immunofluorescence staining (**bottom**) in distal pulmonary arteries from control and SuHx mice treated with saline or Lidd. Scale bar, 25 µm. **F)** Quantification of pulmonary arterial muscularization in distal pulmonary arteries with diameters of 20-100 μm in **E**. Vessels were classified as non-muscularized, partially muscularized, or fully muscularized (n ≥ 5). **G-H**) Quantification of pulmonary arterial remodeling (vessel wall area/total vessel area) in distal pulmonary arteries with diameters of 20-50 µm (**G**) or 50-100 µm (**H**) from H&E-stained sections in **E** (n ≥ 5). **I)** Representative immunofluorescence images of proliferating cell nuclear antigen (PCNA, red) and α-SMA (green) in distal pulmonary arteries. Scale bar, 25 µm. **J)** Quantification of PCNA^+^α-SMA^+^ cells within the α-SMA^+^ population from **I** (n = 5). Data are presented as mean ± SEM. Statistical analyses were performed using two-way ANOVA followed by Tukey’s multiple-comparisons test. Nor, normoxia; SuHx, SU5416/hypoxia; RVSP, right ventricular systolic pressure; RV/LV+S, ratio of right ventricular weight to left ventricular plus septal weight.

### Lidd Suppresses Progression of Monocrotaline -Induced PH and Vascular Remodeling in rats

We then explored the therapeutic effect of Lidd on PH and pulmonary vascular remodeling in monocrotaline (MCT)-treated rats PH model. After 2 weeks, rats from the MCT group were randomly assigned to receive either daily saline or Lidd (50 mg/kg) treatment (**Figure 2A**). As previously reported, MCT challenge significantly elevated RVSP, RV/(LV+S) ratio, and pulmonary vascular wall thickness at 4 weeks (**Figure 2B-H**). Here we found Lidd administration markedly attenuated the increased RVSP (Lidd, 27.53±1.50 mm Hg versus vehicle, 34.71±1.16 mm Hg; P=0.0017; **Figure 2B-2C**), RV/LV+S ratio (Lidd, 0.228±0.023 versus vehicle, 0.348±0.0257; P=0.0024; **Figure 2D**), pulmonary vascular wall thickness (**Figure 2E, 2G-2H**) and muscularization (**Figure 2F**) in MCT-treated rats. Notably, Lidd suppressed PASMC proliferation (**Figure 2I-J**) in monocrotaline-challenged rats. Likewise, Lidd treatment conferred protection against MCT-induced RV remodeling along with downregulated cardiomyocyte hypertrophy and cardiac fibrosis in rats (**Figure S3**). These results show that Lidd treatment alleviates the progression of pulmonary vascular remodeling in rat established pulmonary hypertension.

**Figure 2.**
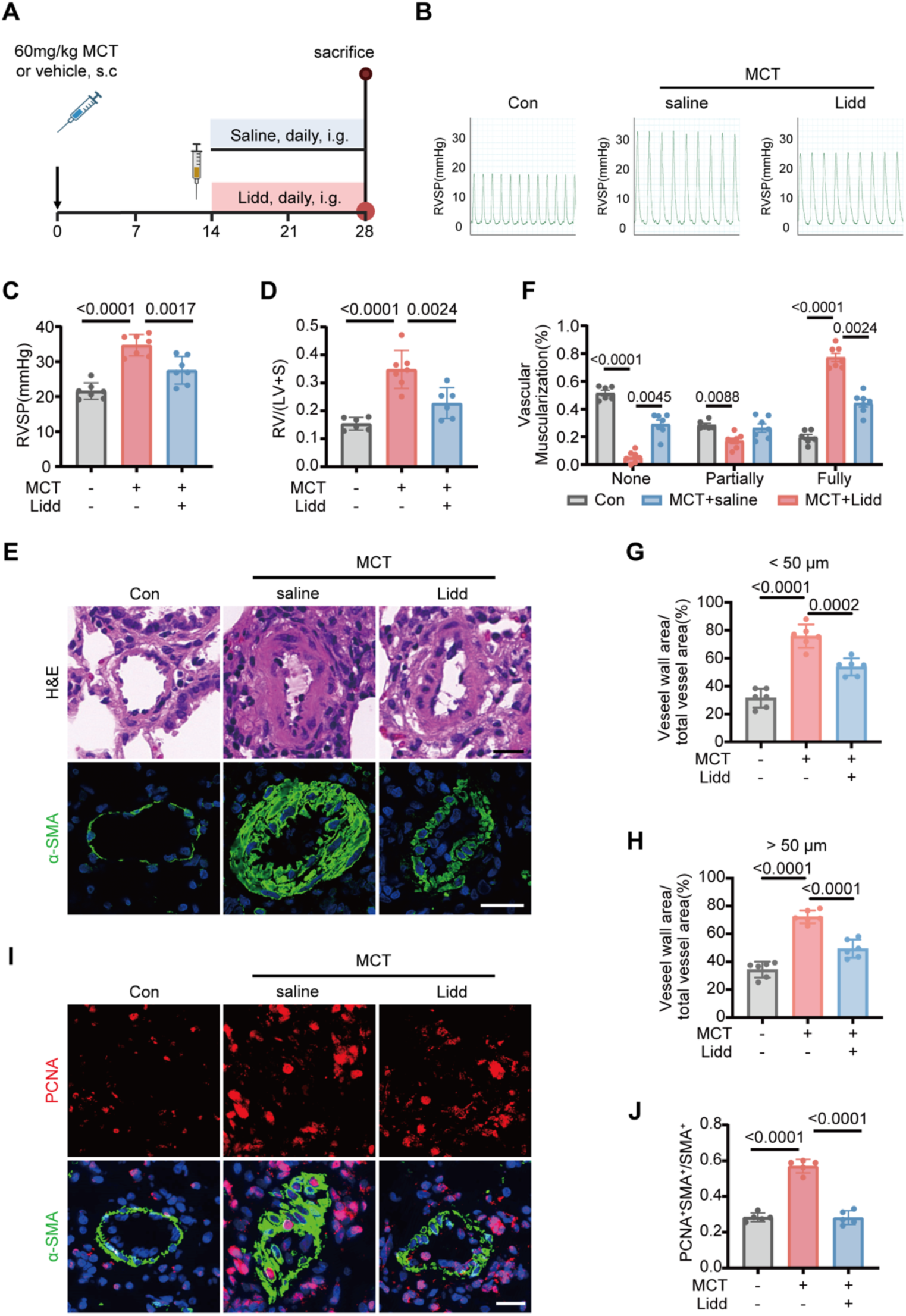
Lidd reverses monocrotaline (MCT)-induced pulmonary hypertension in rat. **A)** Experimental design of the MCT-induced PH mouse model. Mice received a single subcutaneous injection of monocrotaline (MCT; 60 mg/kg) or vehicle on day 0 and were treated with daily oral gavage of Lidd or saline from day 14 to day 28. Hemodynamic measurements and tissue collection were performed on day 28. **B-C**) Representative right ventricular systolic pressure (RVSP) tracings (**B**) and quantification of RVSP (**C**) in control and MCT-treated mice with or without Lidd treatment (n = 7). **D)** Quantification of right ventricular hypertrophy, assessed by the ratio of right ventricular weight to left ventricular plus septal weight (RV/LV+S) (n ≥ 5). **E)** Representative images of hematoxylin and eosin (H&E) staining (**top**) and α-smooth muscle actin (α-SMA, green) immunofluorescence staining (**bottom**) in distal pulmonary arteries from the indicated group. Scale bar, 25 µm. **F)** Quantification of pulmonary arterial muscularization in distal pulmonary arteries with diameters of 20-100 μm in **E**. Vessels were classified as non-muscularized, partially muscularized, or fully muscularized (n ≥ 5). **G - H**) Quantification of pulmonary arterial remodeling (vessel wall area/total vessel area) in distal pulmonary arteries with diameters of 20-50 µm (**G**) or 50-100 µm (**H**) from H&E-stained sections in **E** (n ≥ 5). **I)** Representative immunofluorescence images of proliferating cell nuclear antigen (PCNA, red) and α-SMA (green) in distal pulmonary arteries. Scale bar, 20 µm. **J)** Quantification of proliferating pulmonary arterial smooth muscle cells in **I** (n = 5). Data are presented as mean ± SEM. Statistical analyses were performed using one-way ANOVA followed by Tukey’s multiple-comparisons test. Con, vehicle control (ethanol:saline = 1:4); MCT, monocrotaline; RVSP, right ventricular systolic pressure; RV/LV+S, ratio of right ventricular weight to left ventricular plus septal weight.

### Lidd directly suppresses PASMCs proliferation and inflammatory cytokines expression

To elucidate the molecular mechanisms underlying Lidd-mediated effects on PASMCs, we conducted comprehensive RNA sequencing (RNA-seq) analysis in hPASMCs 24 hours post PDGF-BB and Lidd treatment (**Figure S4A**). RNA-seq analysis identified 508 differentially expressed genes (DEGs; |log2-fold change| >0.58, Q <0.05) post Lidd treatment (**Figure S4B-S4C**), with KEGG analysis showing predominately enrichment of downregulated genes in inflammatory and cytokine-associated pathways, particularly the TNF-α, NF-κB, and IL-17 signaling cascades (**Figure 3A**). Especially, numerous chemokine ligands and inflammatory cytokine receptors were significantly reversed post Lidd treatment (**Figure 3B**). Subsequent q-PCR analysis confirmed the elevated chemokines expression in PDGF-BB induced PASMCs phenotype transition were all attenuated to a low level post Lidd treatment (**Figures 3C**). To confirm the inflammatory phenotypic transition of PASMCs, the co-cultured experiment was designed (**Figures 3D**) and found PDGF-BB-treated PASMCs induced more macrophage transwell and were attenuated by Lidd treatment (**Figures 3E-3F**). Consequently, as seen in vivo, Lidd treatment inhibited PDGF-BB-induced proliferation (**Figure 3G**) and migration (**Figure 3H-J**) of hPASMCs in vitro. To determine whether this immunomodulatory effect also occurs in vivo, we examined perivascular macrophage accumulation in the SuHx-induced mouse and MCT-induced rat pulmonary hypertension models. Lidd treatment markedly reduced perivascular inflammation in both models, as evidenced by decreased CD68⁺ macrophage infiltration (**Figures 3K-N**). Given the emerging recognition of PASMCs immunomodulatory role in shaping the vascular remodeling microenvironment, these findings highlight a direct effect of Lidd in modulating the immune-regulation of PASMC in vascular remodeling.

**Figure 3.**
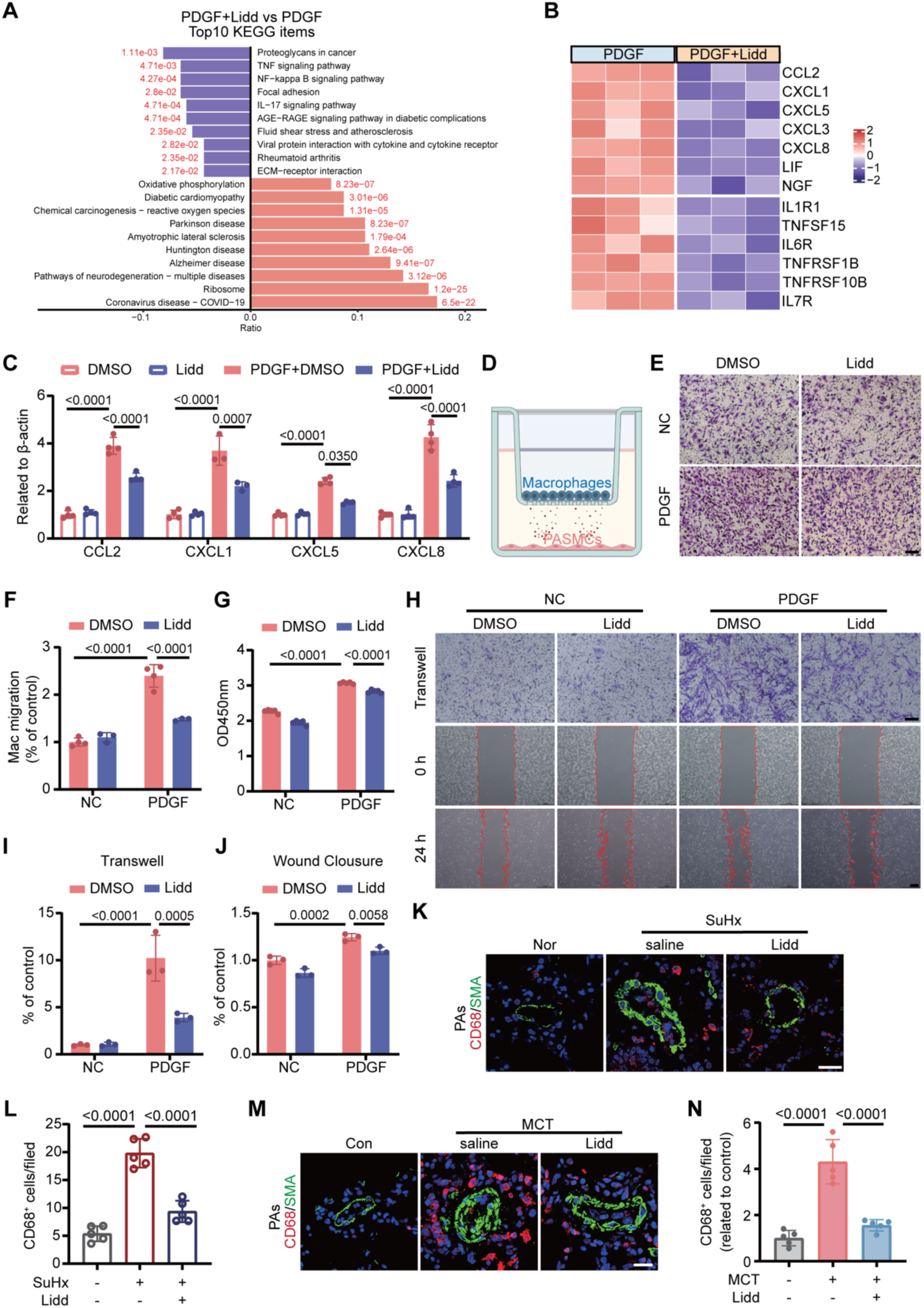
Lidd alleviates pulmonary artery smooth muscle cell (PASMC) inflammation genes expression and phenotypic transformation. hPASMCs were stimulated with PDGF-BB (20 ng/ml) or vehicle and treated with Lidd (100 μM) or DMSO for the indicated times. **A)** Kyoto Encyclopedia of Genes and Genomes (KEGG) enrichment analysis of differentially expressed genes in PDGF-BB-stimulated hPASMCs treated with or without Lidd for 24 h. The top 10 enriched pathways are shown. **B)** Heatmap of representative inflammation-related genes identified by RNA sequencing. **C)** qRT-PCR analysis of CCL2, CXCL1, CXCL5, and CXCL8 mRNA expression in hPASMCs stimulated with PDGF-BB and treated with or without Lidd for 2 h (n = 3). **D)** Schematic of the transwell co-culture system used to examine hPASMC-mediated macrophage recruitment. **E)** Representative images of macrophages migrating toward hPASMCs pretreated with PDGF-BB with or without Lidd for 2 h, followed by 6 h of transwell co-culture. Scale bar, 100 μm. **F)** Quantification of migrated macrophages in **E** (n = 3-4). **G)** Cell proliferation measured by Cell Counting Kit-8 (CCK-8) assay in hPASMCs stimulated with PDGF-BB and treated with or without Lidd for 48 h (n = 3). **H)** Representative images of transwell migration (**top**) and wound healing (**bottom**) assays in hPASMCs stimulated with PDGF-BB and treated with or without Lidd for 24 h. Scale bars, 100 μm. **I-J**) Quantification of transwell migration (**I**) and wound closure (**J**) in **H** (n = 3). **K)** Representative images of immunofluorescence staining for CD68 (red) and α-SMA (green) in PAs from SuHx-exposed mice treated with saline or Lidd. Scale bar, 25 µm. **L)** Quantification of CD68-positive cells in **K**. Cells were counted in five random fields of view (250×250 µm) per sample (n = 5). **M)** Representative images of immunofluorescence staining for CD68 (red) and α-SMA (green) in PAs from MCT-treated mice treated with saline or Lidd. Scale bar, 20 µm. **N)** Quantification of CD68-positive cells in **M**. Cells were counted in five random fields of view (250×250 µm) per sample (n = 5). Data are presented as mean ± SEM. Statistical analyses were performed using one-way (**L**, **N**) or two-way (**C**, **F, G, I, J**) ANOVA followed by Tukey’s multiple-comparisons test. PDGF, platelet-derived growth factor-BB; Lidd, liriodendrin; NC, negative control; DMSO, dimethyl sulfoxide; Nor, normoxia; SuHx, SU5416/hypoxia; Con, vehicle control; MCT, monocrotaline.

### Lidd inhibits PDGF-BB-induced PASMCs phenotypic transformation by reducing glycolytic flux and lactate production

Metabolic reprogramming underlies immunological changes and vascular remodeling^31^. RNA sequencing of PDGF-BB-stimulated hPASMCs revealed the capacity of Lidd to reverse metabolic reprogramming to elevated oxidative phosphorylation (**Figure 3A**). Further GO enrichment analysis showed significant upregulation of oxidative phosphorylation (OXPHOS)-related pathways in Lidd-treated versus PDGF-only groups (**Figure 4A**), counteracting the Warburg-like effect in PH, a metabolic reprogram indicative of bioenergetic insufficiency. Complementary transcriptomic analyses demonstrated coordinated suppression of glycolytic enzymes and induction of oxidative phosphorylation related genes (**Figures S5**), reflecting an inhibition of the pathological metabolic shift toward glucose dependency post PDGF-BB treatment. Consistent with transcriptomic findings, Lidd treatment markedly reduced the glycolytic metabolite -lactate accumulation in SuHx-PH murine lungs (**Figure 4B**) and PDGF-stimulated hPASMC supernatants (**Figure 4C**). Functional assessment via extracellular acidification rate (ECAR) measurements confirmed attenuated glycolytic activity in Lidd-treated cells, with significant reductions in glycosis, glycolylic capacity and glycolylic reserve (**Figure 4D-E**).

**Figure 4.**
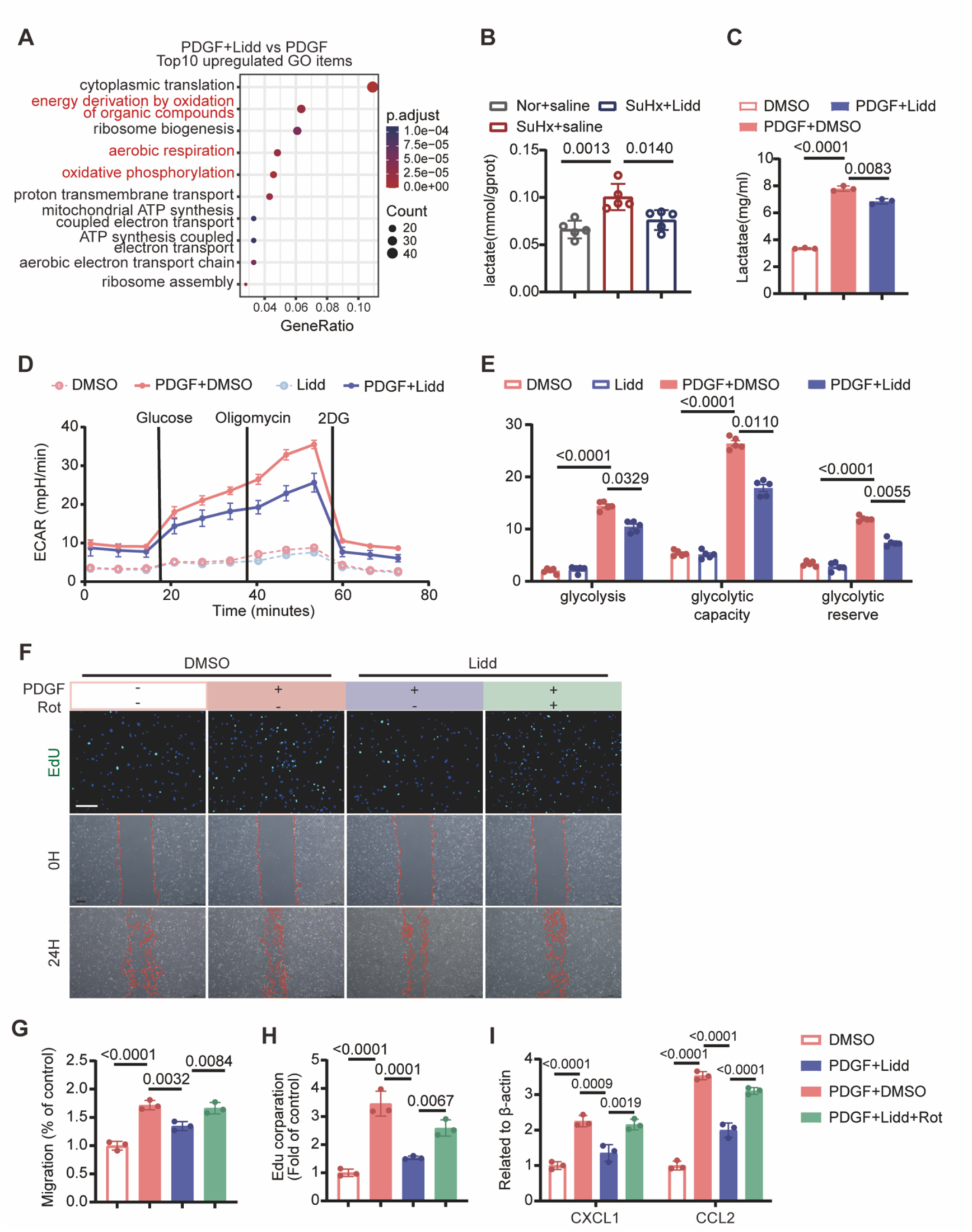
Lidd suppresses PDGF-BB-induced hPASMC phenotypic transformation by reducing glycolysis-derived lactate. hPASMCs were treated with PDGF-BB (20 ng/ml) or vehicle in the presence or absence of Lidd (100 μM) unless otherwise indicated. Rotenone (Rot, 5 nM) was added where indicated. **A)** Gene Ontology (GO) analysis of upregulated genes in Lidd-treated hPASMCs under PDGF-BB stimulation after 24 h of treatment. **B)** Lactate levels in lung tissue from Nor or SuHx-exposed mice treated with saline or Lidd (n = 5). **C)** Lactate levels in the culture supernatant of PDGF-BB-stimulated hPASMCs treated with Lidd or DMSO for 48 h (n = 3). **D)** Representative extracellular acidification rate (ECAR) profiles of control and PDGF-BB-stimulated hPASMCs treated with Lidd or DMSO for 48 h (n = 5). **E)** Quantification of basal glycolysis (**left**), glycolytic capacity (**middle**) and glycolytic reserve (**right**) in **D** (n = 5). **F)** Representative EdU (**top**) and wound healing (**bottom**) assays of hPASMCs under the indicated treatment conditions after 48 h and 24 h of treatment, respectively. Scale bars, 200 µm. **G-H**) Quantification of EdU incorporation (**G**) and wound closure **(H**) in **F** (n = 3). **I)** qRT-PCR analysis of CXCL1 and CCL2 mRNA expression in hPASMCs under the indicated treatment conditions for 2 h (n=3). Data are presented as mean ± SEM. Statistical analyses were performed using one-way (**B**, **C**, **G**, **H** and **I**) or two-way ANOVA (**E**), followed by the Tukey’s multiple-comparisons test. PDGF, platelet-derived growth factor-BB; Nor, normoxia; SuHx, SU5416/hypoxia; ECAR, extracellular acidification rate; DMSO, dimethyl sulfoxide; 2DG, 2-deoxy-D-glucose; EdU, 5-ethynyl-2ʹ-deoxyuridine; Rot, Rotenone; hPASMCs, human pulmonary artery smooth muscle cells.

Previous research has shown that lactate promotes the phenotypic transformation of VSMCs to a synthetic phenotype characterized by increased proliferation and migration. To investigate whether Lidd inhibits PDGF-BB-induced proliferation, migration, and inflammation in hPASMCs by reducing lactate production, exogenous sodium lactate supplement and endogenous mitochondrial respiration inhibitor rotenone were used. Remarkably, exogenous lactate supplementation completely abrogated Lidd mediated anti-proliferative effects and restored migration capacity (**Figure S6**). Similarly, rotenone-induced mitochondrial inhibition abrogated the anti-proliferative and migration effects (**Figure 4F-H**). Notably, the down-regulated mRNA expression levels of the proinflammatory chemokines CXCL1 and CCL2 were also re-upregulated upon mitochondrial inhibition (**Figure 4I**). Thus, Lidd mediated-inhibition of pathological PASMCs phenotypes and inflammatory regulation hinges on the metabolic reprogramming by coordinately enhancing OXPHOS while suppressing glycolysis.

### Lidd blocks PDGF-BB-induced PASMCs phenotype transformation through direct targeting of PFKFB3

PFKFB3, a key glycolytic regulator promoting lactate production, is implicated in pathological PASMCs phenotypic switching in PH^32,33^. Computational molecular docking analysis revealed a strong affinity between Lidd and PFKFB3 (**Figure 5A**). This was functionally validated through drug affinity responsive target stability (DARTS) assays demonstrating Lidd-induced PFKFB3 protein destabilization (**Figure 5B-C**). Notably, Lidd administration reversed PDGF-BB-mediated PFKFB3 upregulation in hPASMCs at both protein expression (**Figure 5D-E**) and functional levels, as evidenced by suppressed PFK enzyme activity (**Figure 5F**). To further confirm the role of PFKFB3 in Lidd-mediated phenotypic transformation of PASMCs, we performed PFKFB3 knockdown using siRNA (**Figure 5G**). Both genetic silencing of PFKFB3 and Lidd administration abolished PDGF-BB-induced proliferative, migratory effects and inflammatory responses (**Figure 5H-K**), but PFKFB3 depletion couldn’t enhance the effects further, confirming that Lidd regulates these processes mainly via targeting PFKFB3.

**Figure 5.**
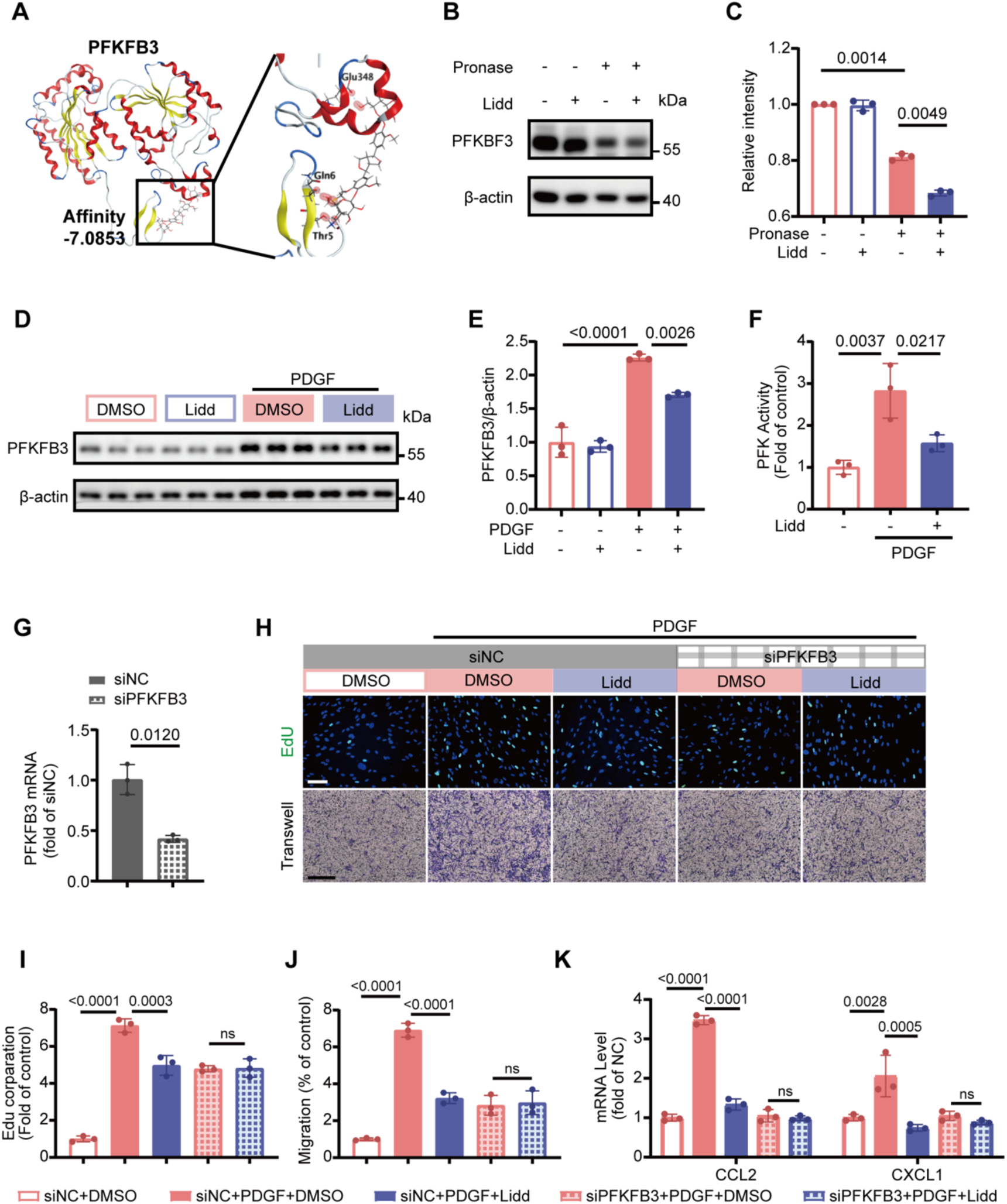
Lidd inhibits PDGF-BB-induced hPASMC phenotypic transformation by directly binding to and reducing 6-Phosphofructo-2-kinase/fructose-2,6-bisphosphatase 3 (PFKFB3). hPASMCs were treated with vehicle or PDGF-BB (20 ng/mL) in the presence or absence of Lidd (100 μM) unless otherwise indicated. siPFKFB3 or control siRNA (siNC) was transfected 24 h before PDGF-BB stimulation where indicated. **A)** Predicted binding mode of Lidd to PFKFB3 determined by molecular docking. **B-C**) Drug affinity responsive target stability (DARTS) assay of hPASMC lysates incubated with Lidd or DMSO followed by pronase digestion. Representative immunoblotting (**B**) and quantification (**C**) of PFKFB3 protein levels are shown (n = 3). **D-E**) Representative immunoblotting (**D**) and quantification (**E**) of PFKFB3 protein levels in hPASMCs under the indicated treatment conditions (n=3). **F)** Intracellular phosphofructokinase-1 (PFK-1) activity in hPASMCs under the indicated treatment conditions (n=3). **G)** qRT-PCR analysis of PFKFB3 mRNA expression in hPASMCs transfected with siPFKFB3 or control siRNA (siNC) (n=3). **H)** Representative EdU assay (**top**) after 48 h of treatment and transwell migration assay (**bottom**) after 24 h of treatment in siNC- or siPFKFB3-transfected hPASMCs under the indicated treatment conditions. Scale bars, 200 μm. **I-J**) Quantification of EdU incorporation (**I**) and transwell migration (**J**) in **H** (n=3). **K**) qRT-PCR analysis of CXCL1 and CCL2 mRNA expression in siNC- or siPFKFB3-transfected hPASMCs under the indicated treatment conditions after 2 h (n=3). Data are presented as mean ± SEM. Statistical analyses were performed using unpaired two-tailed Student’s t-test (**G**), one-way (**B, F, I, J,** and **K**) or two-way (**E**) ANOVA followed by Tukey’s multiple-comparisons test. PFKFB3, 6-phosphofructo-2-kinase/fructose-2,6-bisphosphatase 3; PFK, phosphofructokinase-1; siNC, negative control siRNA.

### Lidd promotes PFKFB3 degradation through FZR1-mediated ubiquitination

To elucidate the molecular mechanism underlying Lidd-mediated PFKFB3 regulation in PASMCs, we systematically evaluated the potential regulatory axes from transcriptional to translational, and post-translational mechanisms. Quantitative PCR analysis revealed preserved PFKFB3 mRNA expression following Lidd treatment in both in vivo models and cultured PASMCs (**Figure 6A-B**). Subsequently, Cycloheximide chase assays demonstrated accelerated PFKFB3 protein degradation with Lidd treatment (**Figure 6C**). Moreover, MG132 (proteasome inhibitor) pretreatment reversed Lidd-induced PFKFB3 reduction (**Figure 6D**). These findings indicated Lidd enhances ubiquitin-dependent proteasomal degradation of PFKFB3. As expected, immunoprecipitation assays in Flag-PFKFB3-transfected 293T cells demonstrated markedly increased polyubiquitination signals in Lidd-treated samples (**Figure 6E**), directly confirming the enhanced ubiquitin conjugation.

**Figure 6.**
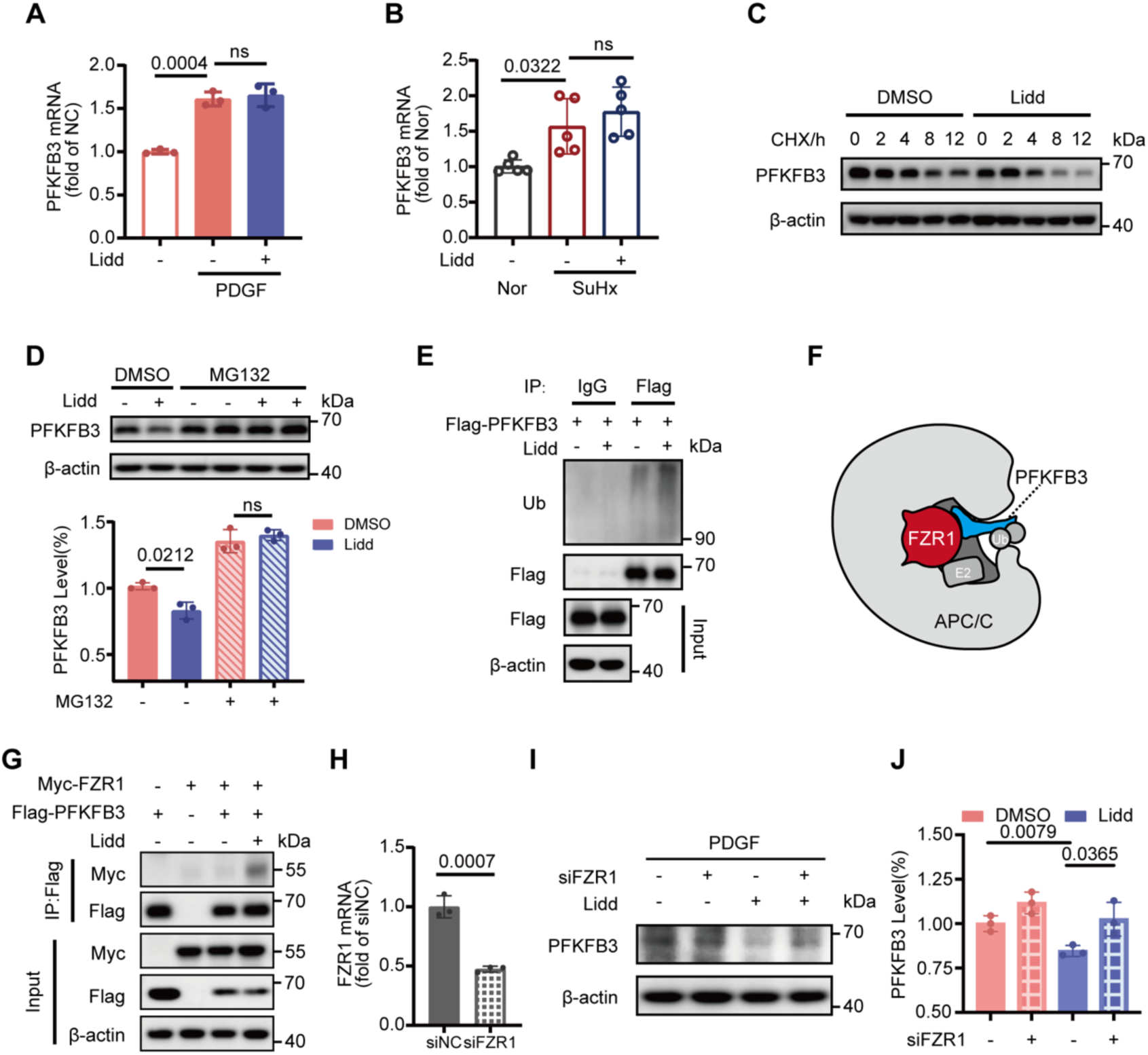
Lidd promotes PFKFB3 binding to Fizzy-related Protein Homolog (FZR1), enhancing its ubiquitination and proteasomal degradation. hPASMCs were treated with PDGF-BB (20 ng/mL) or vehicle in the presence or absence of Lidd (100 μM) unless otherwise indicated. **A)** qRT-PCR analysis of PFKFB3 mRNA levels in hPASMCs under the indicated conditions for 24 h (n = 3). **B)** qRT-PCR analysis of PFKFB3 mRNA levels in lung tissues from Nor or SuHx-exposed mice treated with saline or Lidd (n = 5). **C)** Representative immunoblotting of PFKFB3 protein in PDGF-BB-stimulated hPASMCs treated with DMSO or Lidd for 48 h, followed by cycloheximide (CHX) treatment. Protein lysates were collected at the indicated time points after CHX addition. **D)** Representative immunoblotting (**top**) and quantification (**bottom**) of PFKFB3 protein levels in PDGF-BB-stimulated hPASMCs treated with DMSO or Lidd for 48 h followed by MG132 (100 µM) treatment for the final 6 h where indicated (n = 3). **E)** Representative anti-Flag immunoprecipitation followed by immunoblot analysis for ubiquitin (Ub) and Flag in 293T cells transfected with Flag-PFKFB3 and treated with DMSO or Lidd for 48 h. **F)** Schematic illustrating FZR1-mediated ubiquitination of PFKFB3 through the anaphase-promoting complex/cyclosome (APC/C) E3 ubiquitin ligase complex. **G)** Representative anti-Flag immunoprecipitation followed by immunoblot analysis for Myc and Flag in 293T cells co-transfected with Myc-FZR1 and Flag-PFKFB3 and treated with DMSO or Lidd for 48 h. **H)** qRT-PCR analysis of FZR1 mRNA expression in hPASMCs transfected with siFZR1 or siNC (n = 3). **I-J**) Representative immunoblotting **(I)** and quantification **(J)** of PFKFB3 protein levels in PDGF-BB-stimulated hPASMCs treated with DMSO or Lidd following transfection with siFZR1 or siNC (n = 3). Data are presented as mean ± SEM. Statistical analyses were performed using an unpaired two-tailed Student’s t-test (**H**), one-way ANOVA (**A** and **B**), or two-way ANOVA (**D** and **J**), followed by Tukey’s multiple-comparisons test. CHX, cycloheximide; Ub, ubiquitination; FZR1, Fizzy-Related Protein Homolog; APC/C, anaphase-promoting complex/cyclosome; siNC, negative control siRNA.

Previous studies indicate that PFKFB3 binds to ubiquitin ligase APC/C via FZR1 and undergoes proteasomal degradation following ubiquitination^34,35^ (**Figure 6F**). Co-immunoprecipitation analyses revealed strengthened PFKFB3-FZR1 complex formation upon Lidd treatment (**Figure 6G**). Crucially, genetic ablation of FZR1 completely abolished both Lidd-induced ubiquitination and degradation of PFKFB3 (**Figure 6H-J**). These data indicate Lidd acts as a molecular enhancer of the PFKFB3-FZR1 interaction, thereby potentiating APC/C-mediated ubiquitination and subsequent proteasomal degradation.

### Lidd inhibits PASMCs phenotypic transition by inhibiting H3K18 lactylation-mediated transcriptional activation of CCND1, TNC, and CCL2

Histone lactylation, a recently identified metabolite-sensitive epigenetic mark, bridges glycolytic flux with transcriptional reprogramming in PH pathogenesis^36–39^. Given the established role of lactate in promoting synthetic VSMC phenotypes through H3K18 lactylation^37,40^, we tested whether Lidd suppresses PDGF-BB-induced PASMC phenotype transformation by disrupting H3K18 lactylation-mediated transcriptional reprogramming. Immunoblotting showed robust PDGF-BB-induced global histone lactylation in PASMCs, an effect markedly attenuated by Lidd pretreatment (**Figure 7A**). Targeted assays confirmed Lidd significantly inhibited PDGF-BB-induced H3K18 lactylation (**Figure 7B**). Conversely, mitochondrial complex I inhibition (rotenone) or exogenous lactate supplementation (NaLa) re-amplified Lidd attenuated H3K18 lactylation (**Figure 7C-D**, Figure S7). P300 was identified as a critical histone lactyltransferase^41,42^, and P300 knockdown showed consistent effects with Lidd-mediated attenuation of H3K18 lactylation (**Figures 7E-F**), paralleled with diminished PASMC proliferation (**Figure 7G-7H**) and migration (**Figure 7I-J**) upon rotenone treatment. Notably, data analysis of the published results^43^ revealed that H3K18la enrichment at gene promoters is strongly correlated with gene transcription, including the three PH–relevant targets: the cell cycle regulator CCND1, the extracellular matrix remodeler TNC, and the proinflammatory chemokine CCL2 (**Figure S9A-S9C**). Consistent with these findings, ChIP-qPCR validated the increased H3K18la enrichment at the Ccnd1, Tnc, and Ccl2 promoter regions under PDGF-BB stimulation, which were suppressed in Lidd treatment group (**Figures 7K-M**). Correspondingly, Lidd suppressed PDGF-BB-induced upregulation of these genes, while rotenone-induced lactate accumulation restored their expression (**Figures S8D-F**). Collectively, our results identify a lactate–H3K18la–CCND1/TNC/CCL2 axis through which Lidd restrains PASMC proliferative and inflammatory phenotypic transition, consequently alleviating vascular remodeling and vascular inflammation **(Figure S10**), offering a druggable pathway for pulmonary vascular remodeling.

**Figure 7.**
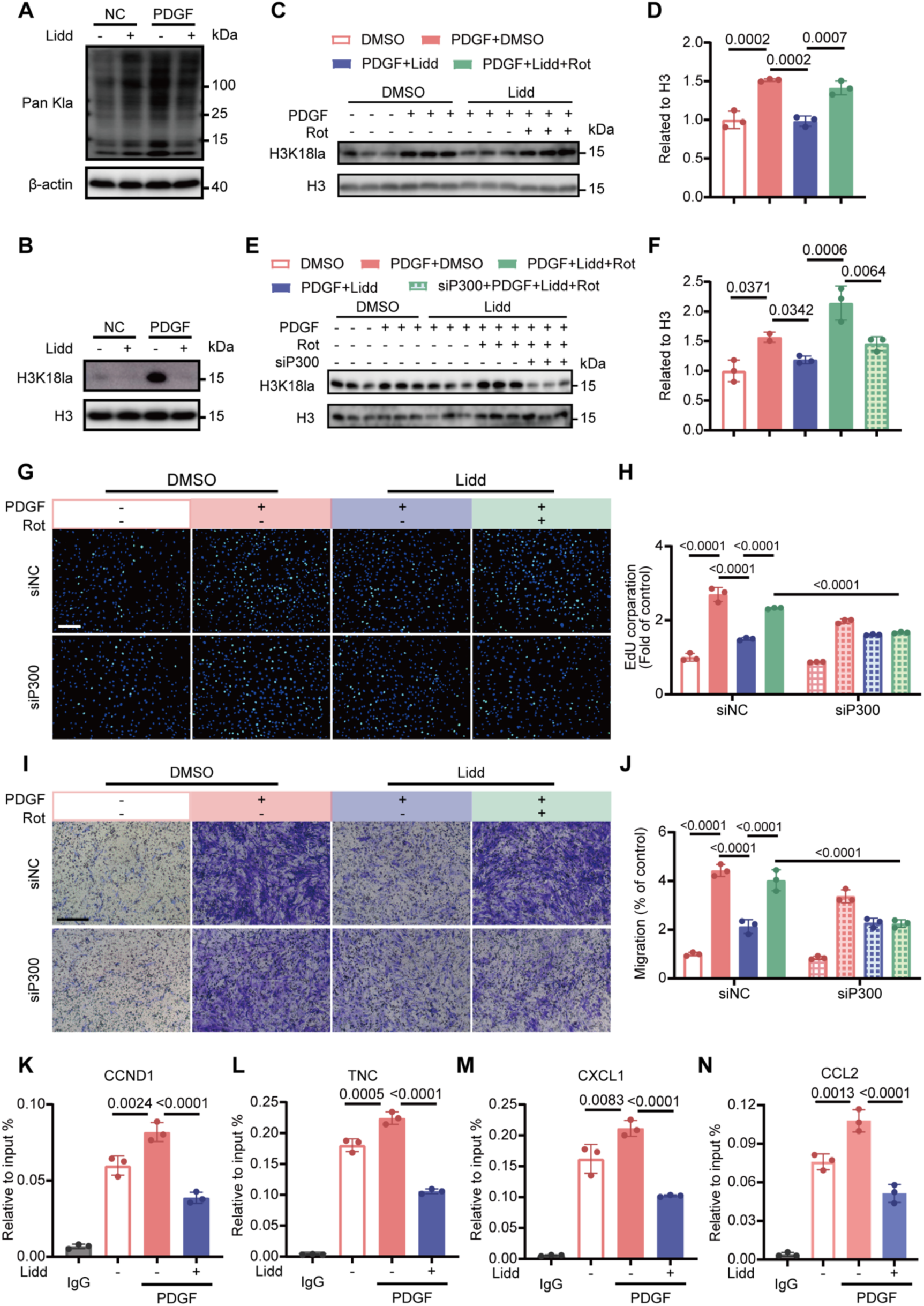
Lidd prevents PDGF-BB-induced PASMC phenotypic transformation by inhibiting H3K18 lactylation-mediated activation of CCND1, TNC, and CCL2 transcription. hPASMCs were treated with vehicle or PDGF-BB (20 ng/mL) in the presence or absence of Lidd (100 µM). Rotenone (Rot, 5nM) was added where indicated. **A)** Representative immunoblotting of pan-lysine lactylation (Pan Kla) in hPASMCs under the indicated treatment conditions. **B)** Representative immunoblotting of histone H3 lysine 18 lactylation (H3K18la) in hPASMCs under the indicated treatment conditions. **C-D**) Representative immunoblotting (**C**) and quantification (**D**) of H3K18la levels in hPASMCs under the indicated treatment conditions (n = 3). **E-F**) Representative immunoblotting (**E**) and quantification (**F**) of H3K18la levels in siNC- or siP300-transfected hPASMCs under the indicated treatment conditions (n = 3). **G-H**) Representative EdU assay (**G**) and quantification of EdU incorporation (**H**) in siNC- or siP300-transfected hPASMCs under the indicated treatment conditions. Scale bar, 200 μm (n = 3). **I-J**) Representative transwell migration assay (**I**) and quantification of migrated cells (**J**) in siNC- or siP300-transfected hPASMCs under the indicated treatment conditions. Scale bar, 200 μm (n = 3). **K-N**) ChIP-qPCR analysis of H3K18la enrichment at the promoters of CCND1 (**K**), TNC (**L**), CXCL1(**M**), and CCL2 (**N**) in hPASMCs under the indicated treatment conditions (n = 3). Data are presented as mean ± SEM. Statistical analyses were performed using one-way (**K-N**) or two-way (**D**, **F**, **H**, and **J**) ANOVA followed by Tukey’s multiple-comparisons test. Rot, rotenone; Pan Kla, pan-lysine lactylation; H3K18la, histone H3 lysine 18 lactylation; H3, histone H3; siNC, negative control siRNA.

### PFKFB3 is essential for the Lidd–mediated alleviation of pulmonary hypertension in vivo

To further investigate whether PFKFB3 is required for the therapeutic efficacy of Lidd in PH, we evaluated the impact of Lidd treatment in SuHx-induced PH using PFKFB3^+/−^ and wild-type littermates (**Figure 8A**). As expected, PFKFB3 haploinsufficiency alone significantly improved pulmonary hemodynamics, indicated by a reduction in RVSP, RV/(LV+S), and an increase in the ratio of PA AT/ET, TAPSE (**Figure 8B-D**), along with attenuation of pulmonary vascular remodeling, including decreased medial wall thickness, reduced muscularization and PASMC proliferation (**Figure 8E-I**), as well as attenuated perivascular inflammation (**Figure 8J-K**). In addition, PFKFB3 haploinsufficiency attenuated right ventricular remodeling, as demonstrated by reduced cardiomyocyte hypertrophy and collagen deposition (**Figure S8A-C**). Importantly, Lidd failed to provide any further therapeutic benefit in PFKFB3^+/−^ mice. Measures of pulmonary hemodynamics, arterial wall remodeling, inflammatory cell infiltration, and right ventricular structural changes remained comparable between saline- and Lidd-treated PFKFB3^+/−^ mice (**Figure 8B-K** and **Figure S11**). Taken together, these results indicate that PFKFB3 is a critical target required for Lidd’s protective actions against PH, and that inhibition of PFKFB3-mediated glycolysis constitutes the key mechanism through which Lidd ameliorates pulmonary vascular remodeling and disease progression.

**Figure 8.**
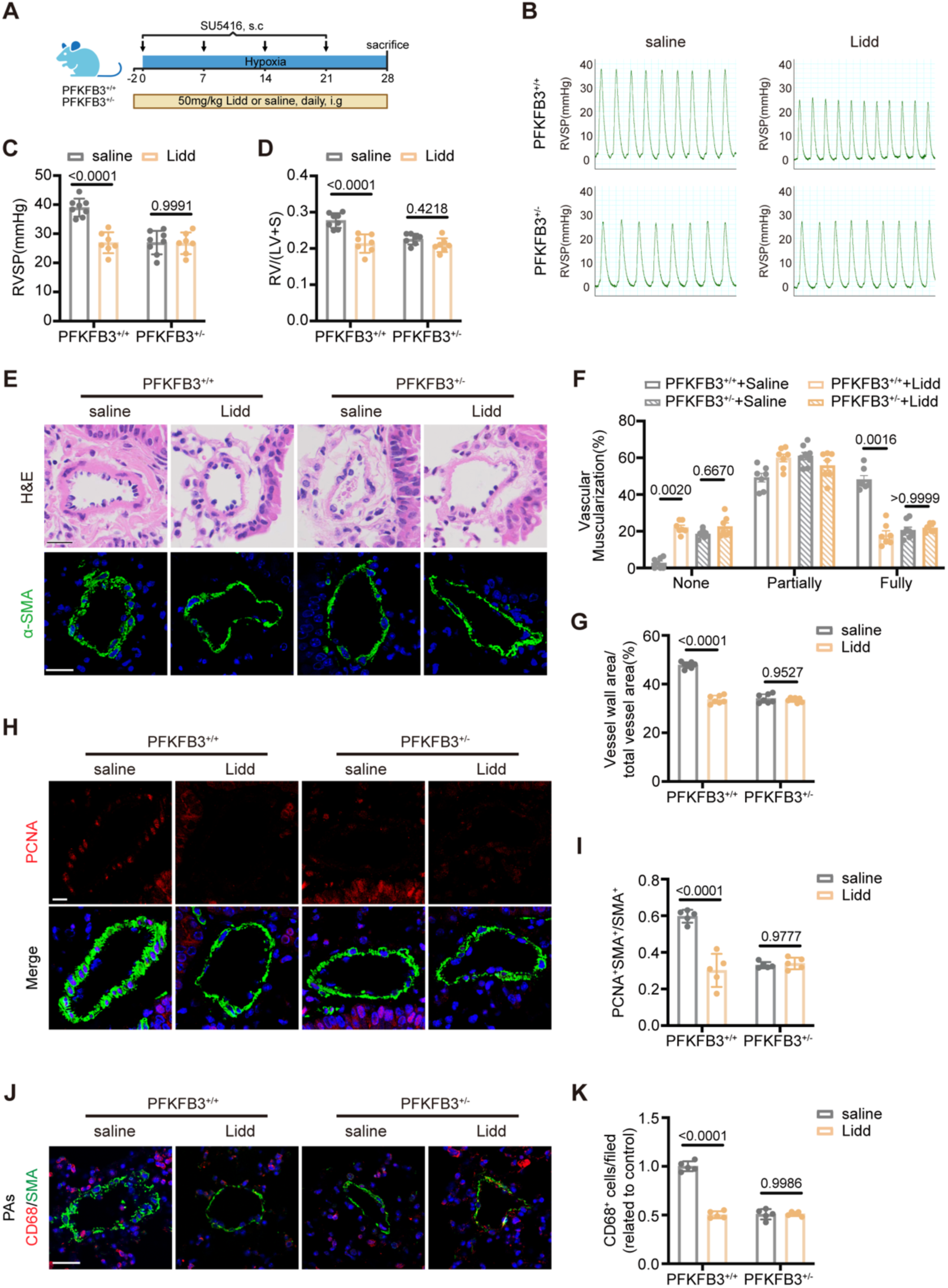
Lidd fails to provide additive protection against SuHx-induced pulmonary hypertension in PFKFB3-deficient mice. **A)** Experimental design of the SuHx-induced PH model and Lidd treatment in PFKFB3^+/+^ and PFKFB3^+/-^ mice. **B-C)** Representative right ventricular systolic pressure (RVSP) tracings (**B**) and quantification of RVSP (**C**) in SuHx-exposed PFKFB3^+/+^ and PFKFB3^+/-^ mice treated with saline or Lidd (n = 7). **D)** Quantification of right ventricular hypertrophy, assessed by the ratio of right ventricular weight to left ventricular plus septal weight (RV/LV+S) in SuHx-exposed PFKFB3+/+ and PFKFB3+/- mice treated with saline or Lidd (n = 7). **E)** Representative images of hematoxylin and eosin (H&E) staining (**top**) and α-smooth muscle actin (α-SMA, green) immunofluorescence staining (**bottom**) in distal pulmonary arteries from SuHx-exposed PFKFB3^+/+^ and PFKFB3^+/-^ mice treated with saline or Lidd. Scale bar, 25 µm. **F)** Quantification of pulmonary arterial muscularization in distal pulmonary arteries with diameters of 20-100 μm in **E**. Vessels were classified as non-muscularized, partially muscularized, or fully muscularized (n ≥ 5). **G)** Quantification of pulmonary arterial remodeling (vessel wall area/total vessel area) in distal pulmonary arteries with diameters of 20-100 µm from H&E-stained sections in **E** (n ≥ 5). **H)** Representative immunofluorescence images of proliferating cell nuclear antigen (PCNA, red) and α-SMA (green) in distal pulmonary arteries. Scale bar, 20 µm. **I)** Quantification of PCNA^+^α-SMA^+^ cells within the α-SMA^+^ population from **H** (n = 5). **J)** Representative images of immunofluorescence staining for CD68 (red) and α-SMA (green) in PAs from SuHx-exposed PFKFB3^+/+^ and PFKFB3^+/-^ mice treated with saline or Lidd. Scale bar, 25 µm. **K)** Quantification of CD68-positive cells in **J**. Cells were counted in five random fields of view (250×250 µm) per sample (n = 5). Data are presented as mean ± SEM. Statistical analyses were performed using two-way ANOVA followed by Tukey’s multiple-comparisons test. Nor, normoxia; SuHx, SU5416/hypoxia; RVSP, right ventricular systolic pressure; RV/LV+S, ratio of right ventricular weight to left ventricular plus septal weight; PCNA, proliferating cell nuclear antigen; α-SMA, α-smooth muscle actin.

## Discussion

Our study establishes Liriodendrin (Lidd) as a dual-action (anti PASMC proliferation and inflammatory phenotypic transition) therapeutic agent against vascular remodeling in pulmonary hypertension (PH), operating through PFKFB3-mediated glycolytic-H3K18la regulation. In both SuHx and MCT models, Lidd attenuated PH progression by concurrently suppressing vascular inflammation and remodeling. Mechanistically, Lidd binds PFKFB3 to induce FZR1-dependent ubiquitination and proteasomal degradation, thereby reducing glycolytic flux and lactate production. This metabolic reprogramming directly diminished histone H3K18 lactylation levels, leading to transcriptional downregulation of key inflammatory (CCL2) and remodeling mediators (CCND1, TNC) in pathological PASMCs remodeling. These findings position Lidd as a novel compound that disrupts the PFKFB3/lactylation axis, offering a promising therapeutic strategy for clinical PH management.

The inflammatory microenvironment is increasingly recognized as a central driver of PH progression^5,44^. While traditional immunotherapies targeting single pathways (e.g., IL-6 or B-cell depletion) show limited clinical efficacy^14,15^, our work reveals PASMCs as active architects of inflammatory signaling. Under PDGF stimulation, PASMCs secrete CXCL1/3/5, CCL2/5, and ICAM1^45–47^, establishing an autonomous inflammatory circuit that sustain microenvironmental activation. This mechanistic transformation is further evidenced by PH-specific disruption of structural-immune communication networks, where inflammatory signaling hubs predominantly localize to vascular structural cells^20,48^. Crucially, PASMCs exhibit coordinated upregulation of cytokine receptors (IL-6R, IL-8R) and ligands (CCL2, CXCL12), transitioning from passive responders to active orchestrators of vascular remodeling^20,46,49,50^. Here we found, Lidd uniquely disrupts this pathogenic axis through dual modulation of PASMC plasticity and innate immunity. With predominately enrichment of downregulated genes in inflammatory and cytokine-associated pathways, particularly the TNF-α, NF-κB, and IL-17 signaling cascades, numerous chemokine ligands and inflammatory cytokine receptors were significantly reversed post Lidd treatment. Moreover, the co-cultured experiment revealed the capacity of Lidd to attenuate PDGF-BB-induced PASMC-macrophage crosstalk, reducing chemotactic migration by 40%. All above demonstrate a superior therapeutic potential of the novel compound Lidd compared to conventional vasodilatory or single-pathway immunotherapies in PH treatment.

Metabolic alterations, particularly heightened glycolysis, critically drive vascular remodeling in PH by fueling cell proliferation, migration, and inflammation^51–54^. PFKFB3, a key glycolytic enzyme, catalyzes the production of fructose-2,6-bisphosphate (F-2,6-P), a potent activator of PFK-1, sustaining glycolytic flux and supporting the pathological remodeling of pulmonary arteries. This PFKFB3-mediated glycolytic surge has been strongly implicated in promoting phenotypic switching of PASMCs in PH^33,53^. Notably, PFKFB3-driven lactate accumulation activates calpain-2/ERK1/2 signaling to promote PASMC proliferation and collagen synthesis^33^. Beyond glycolysis, PFKFB3 regulates multiple signaling pathways, including Cdk1, PI3K/Akt, and NF-κB, underscoring its broad roles in cell cycle progression and inflammation^55^. Unlike conventional PFKFB3 inhibitors (e.g., 3PO) that merely block enzymatic pockets, Lidd induces FZR1-mediated PFKFB3 degradation, disrupting both catalytic and scaffolding functions of PFKFB3, achieving enhanced efficacy at reduced concentrations. While our current study primarily focuses on the direct suppression of Lidd on PASMCs hyperproliferation, migration, and pro-inflammatory phenotypes, the pan-cellular role of PFKFB3 in PH, including endothelial pro-angiogenic cytokine release^53^ and macrophage-mediated vascular inflammation^56^, warrant investigation into Lidd-mediated multicellular effects.

In pulmonary hypertension, pathological hyperactivation of glycolysis in PASMCs drives excessive lactate production and accumulation, thereby linking metabolic dysregulation to pulmonary vascular remodeling^33,51^. This metabolic reprogramming evolves beyond an adaptive response into a pathological driver of vascular remodeling, orchestrated by metabolism-epigenetic crosstalk. Functioning as both metabolic byproduct and epigenetic modulator, lactate induces histone lactylation, a novel post-translational modification regulating chromatin dynamics^36^, has been increasingly implicated in vascular pathology^39,40,43,57,58^. Our study identified pronounced enrichment of lactate-derived H3K18la at transcription start sites of CCND1, TNC, CXCL1 and CCL2 in phenotypically switched VSMCs. Critically, Lidd treatment suppressed transcriptional activation of these genes by inhibiting histone lactylation, an effect reversible upon lactate supplementation, OXPHOS inhibition, or P300 knockdown. This regulatory pattern aligns with established mechanisms linking histone lactylation to multiple signaling pathways related to pulmonary vascular remodeling. For example, elevated intracellular lactate in PASMCs enhances histone lactylation at hypoxia-inducible factor 1 α (HIF-1 α) targets (e.g., Bmp5, Trpc5, Kit), directly driving phenotypic transformation and arterial remodeling^39^. Additionally, lactate-mediated H3K18la promotes METTL3 expression^59^, which interconnects with the METTL3/YTHDF2/PTEN axis to activate PI3K/AKT signaling and exacerbate PASMC proliferation^60^. Beyond cell-autonomous effects, myofibroblast-derived lactate induces histone lactylation in profibrotic gene promoters of macrophages^61^, potentially exacerbating PAH progression through fibrotic crosstalk^62^.These reinforce the emerging role of histone lactylation as a critical bridge linking metabolic signaling to epigenetic regulation in PH treatment.

In conclusion, Lidd represents a promising therapeutic candidate for PH treatment. By degrading PFKFB3 to disrupt the lactate-lactylation axis, Lidd mediates lactylation regulation in vascular smooth muscle cells to disrupt the pathological processes of vascular remodeling and immune-regulation.

### Experimental Section

#### Animals

Eight-to-ten-week-old male mice maintained on a C57BL/6 genetic background and Eight-week-old male Sprague-Dawley rats were purchased from Charles River Laboratories, Beijing. All animals were housed in a specific pathogen-free facility with controlled temperature (22±1°C) and relative humidity (50±5%) on a 12-hour light/dark cycle, with free access to sterile food and water. All animal experiments procedures were performed in accordance with the Animal Care and Use Committees of the Shanghai General Hospital, Shanghai Jiaotong University School of Medicine.

#### Rodent model of PH

To establish a SU5416/hypoxia-induced PH mouse model, mice received subcutaneous injection of SU5416 (20mg/kg) once weekly, and were exposed to hypoxia (10% O_2_) for 4 weeks as previously described^63^. Lidd was administered daily by gavage (50mg/kg) three days before the SuHx exposure to assess its preventive effects. For the monocrotaline-induced rat PH model, rats were subcutaneously injected with MCT (60mg/kg) once and maintained for four weeks. 2 weeks after injection, rats were treated with saline or Lidd (50mg/kg/day) by gavage for another two weeks.

#### Hemodynamic measurement and tissue collection

At the end of treatment, hemodynamic profiles were assessed as previously described^64^. Animals were anesthetized with pentobarbital (50mg/kg, intraperitoneal injection), then were mechanically ventilated through a transtracheal catheter. Immediately, a 1.2-F (for mice) or 1.4-F (for rats) micro-tip pressure transducer catheter (Millar Instruments, Houston, TX, USA) was carefully inserted into the right ventricle, and RVSP was continuously recorded for 2 min using the PowerLab Data Acquisition System (ADInstruments, Sydney, Australia). Animals with HR < 250/min were excluded a priori from the analysis. After hemodynamic measurements, the pulmonary circulatory system was flushed with chilled PBS, and the heart and lung tissues were collected. The lower lobes of right lungs were fixed in 4% paraformaldehyde (PFA), and left lungs were flash-frozen in liquid nitrogen. The free wall of right ventricle (RV) and left ventricle (LV) plus septum were dissected and weighed. Right ventricular hypertrophy was quantified by measuring the mass ratio of the right ventricular free wall to the LV and septum (RV/(LV+S)). All treatments and analyses were performed in a blinded fashion manner.

#### Pulmonary vascular remodeling analysis

After fixation, the lungs were embedded in paraffin and sectioned at 5 μm thickness, these sections were then stained with hematoxylin and eosin (H&E) for pulmonary vascular remodeling analyses. In each rodent, a total of 20-30 intra-acinar vessels accompanying either alveolar ducts or alveoli were examined by an observer who was blinded to the treatment. Each vessel was categorized into non-muscularized, partially muscularized, and fully muscularized. The proportion of muscularized pulmonary vessels was calculated as dividing the number of partially or fully muscular vessels by the total number counted in the same sample. The medial wall thickness of intra-acinar arteries was calculated with Fiji software. The medial wall thickness was expressed as the ratio of the medial area to the entire vessel area.

#### Histological assessment of hearts

The RV was fixed in 4% PFA, embedded in paraffin, and sectioned at 5 μm thickness. Paraffin sections were stained with H&E and Sirius red. RV cardiomyocyte size was determined by measuring cardiomyocyte’s cross-sectional area. Degree of RV fibrosis was evaluated as the ratio of red-colored area to the total area in Sirius red staining images at 20X magnification.

#### RNA sequencing and analysis

Total RNA was extracted with the Trizol Reagent (Invitrogen Life Technologies) from mouse lungs or hPASMCs, after which the concentration, quality and integrity were determined using a NanoDrop spectrophotometer (Thermo Scientific). Three micrograms of RNA were used as input material for the RNA sample preparations. Sequencing libraries were generated according to the following steps. Firstly, mRNA was purified from total RNA using poly-T oligo-attached magnetic beads. Fragmentation was carried out using divalent cations under elevated temperature in an Illumina proprietary fragmentation buffer. First strand cDNA was synthesized using random oligonucleotides and Super Script II. Second strand cDNA synthesis was subsequently performed using DNA Polymerase I and RNase H. Remaining overhangs were converted into blunt ends via exonuclease/polymerase activities and the enzymes were removed. After adenylation of the 3′ends of the DNA fragments, Illumina PE adapter oligonucleotides were ligated to prepare for hybridization. To select cDNA fragments of the preferred 400-500 bp in length, the library fragments were purified using the AMPure XP system (Beckman Coulter, Beverly, CA, USA). DNA fragments with ligated adaptor molecules on both ends were selectively enriched using Illumina PCR Primer Cocktail in a 15 cycle PCR reaction. Products were purified (AMPure XP system) and quantified using the Agilent high sensitivity DNA assay on a Bioanalyzer 2100 system (Agilent). The sequencing library was then sequenced on NovaSeq 6000 platform (Illumina) Shanghai Personal Biotechnology Cp. Ltd.

Differential expressions were analyzed using DESeq2. |log2FoldChange|>0.58, *Q*-value < 0.05 was set as the threshold for significantly differentially expressed genes (DEGs). We mapped all the genes to Terms in the Gene Ontology database and calculated the numbers of differentially enriched genes in each Term. Using topGO (v2.50.0) to perform GO enrichment analysis on the differential genes, calculate *p*-value by hypergeometric distribution method (the standard of significant enrichment is *p*-value <0.05), and find the GO term with significantly enriched differential genes to determine the main biological functions performed by differential genes. ClusterProfiler (v4.6.0) software was used to carry out the enrichment analysis of the KEGG pathway of differential genes, focusing on the significant enrichment pathway with *Q*-value <0.05.

#### Immunofluorescence analysis

To explore the infiltration of macrophages, mouse lung sections co-stained with CD68 and α-SMA were performed using a routine double immunofluorescence. Briefly, sections were deparaffinized, antigen retrieved, permeabilized, and nonspecific sites blocked. After that, sections were incubated with primary antibodies. After washing with PBS for 3 times, sections were then incubated with Alexa Fluor-488 or -555 conjugated secondary antibody and counterstained with 4’,6-diamidino-2-phenylindole (DAPI). The stained slides were visualized using confocal fluorescence microscope (Leica TCS SP8 X). Six fields (at 40X magnification) per mouse lung were randomly selected for statistical analysis. The mean of CD68 positive cell numbers in six fields was calculated. To reveal the vascular smooth muscle cells phenotype transformation in vivo, mouse lung sections were co-staining with PCNA and α-SMA, six vessels per mouse lung were assessed. To analyze the result of PCNA and α-SMA co-stained, the Fiji software was used for colocalization analysis.

#### Cell culture and cell treatments

Primary human pulmonary arterial smooth muscle cells (hPASMCs, catalog no. FH-H119) were purchased from FuHeng Biology (Shanghai, China). Primary hPASMCs were cultured in DMEM/F-12 (Gibco, Grand Island, NY, USA) supplemented with 10% fetal bovine serum and 1% penicillin-streptomycin, and were used at passages 3-6. 293T cells were cultured in DMEM. Cells were plated on 6-well culture dishes at a density of 1 X 105 cells per well. For the hypoxia experiments, hPASMCs were placed in a modular incubator chamber with 1% O_2_ for 24 hours. For PDGF-BB (20 ng/ml) treatment, hPASMCs were serum-starved for 24 hours and then treated with PDGF-BB for 24-48 hours. 100 nM Lidd were used to treat PASMCs if it was not indicated. To investigate the mechanism that Lidd suppresses PASMCs phenotype transformation via PFKFB3, the PFKFB3 siRNA was used under PDGF-BB treatment condition. To explore the mechanism of PFKFB3 downregulation, the plasmids of Myc-FZR1 and Flag-PFKFB3 were used in 293T cells. To determine whether Lidd suppresses PASMCs phenotype transformation through decreased lactate production, rotenone (5 nM) or sodium lactate (5 mM) was administrated in culture medium 12 hours after PDGF-BB and Lidd treatment, and incubated for another 24 hours. siRNA sequences were listed in **Table S3.**

#### Cell Counting Kit-8 assay

Cell Counting Kit-8 (Abclonal) was used to determine cell proliferation according to the manufacturer’s instructions. Approximately, 8×10^3^ hPASMCs were seeded in 48-well plates and cultured with the indicated treatments for the indicated times. After each treatment, the medium was removed and replaced with 100 μL fresh medium and 10 μL of Cell Counting Kit-8 solution according to the manufacturer’s instructions. The plates were incubated for 3 hours, and absorbance was measured at 450 nm using a microplate reader.

#### 5-Ethynyl-2’-deoxyuridine (EdU) staining

EdU staining was performed using an EdU Cell Proliferation Assay Kit (Beyotime Biotechnology, China) according to the manufacturer’s protocol. Briefly, hPASMCs were seeded at a density of 2×10^4^ cells/well into 24-well plates, followed by incubation with indicated reagent for the indicated period. After experimental stimulation, cells were incubated with 10 mM EdU for 4 hours before fixation with 4% PFA. The experiment was performed according to the manufacturer’s protocol. 5 frames were examined per well using a Leica microscope (Leica, Heidelberg, Germany), and the relative EdU-positive ratio was calculated with Fiji Software.

#### Wound healing assay

Cells were grown to confluence in 6-well plates and serum starved for 24 hours. A linear scratch was created in the cell monolayer using a 200 μL pipette tip. The cells were subsequently treated with PDGF-BB and Lidd in DMEM-F12 supplemented with 1% fetal bovine serum. Wound closure was assessed by capturing images 24 hours after the scratch. The degree of wound healing was quantified using Fiji software.

#### Transwell assay

Following treatment with the specified reagents, hPASMCs (1×10⁴ cells per 200 μL) were plated in the upper chamber containing serum-free DMEM-F12. The lower chamber was filled with 500 μL of DMEM-F12 supplemented with 20% fetal bovine serum. After 24 hours, the filter was fixed and stained with crystal violet for 15 minutes. The membranes were then washed with distilled water, and non-migrated cells on the upper surface were carefully scraped off. Five random fields of view were selected under the microscope, and images were captured for analysis. Cell migration was quantified by counting the number of cells in each field using Fiji software.

#### RNA extraction and Real-Time Quantitative PCR

Total RNA was isolated from lung homogenates or hPASMCs using TRIzol reagent (Invitrogen, Carlsbad, CA). RNA samples (1.0 μg) were reverse-transcribed to cDNA using the reverse transcription reagent kit (Vazyme, Nanjing, China), according to manufacturer’s instructions. The resulting cDNA was amplified by real-time PCR in a 10-μL reaction volume with SYBR Green Universal Master Mix. All the test measurements were normalized to the mean of the control measurements. The primer sequences used are listed in **Table S4**.

#### Phosphofructokinase (PFK) activity

After various treatment, hPASMCs were collected. Intracellular total PFK1 activity was measured using the PFK Activity Colorimetric Assay Kit (Solarbio, Beijing) following the manufacturer’s instructions. ***Seahorse assay***: ECAR was measured using the Seahorse XFe96 Extracellular Flux Analyzer (Agilent, Billerica, USA), according to the manufacturer’s protocol. Briefly, cells were plated at 5,000 cells per well in XF96 Cell Culture Microplates. The assay was performed in XF base medium supplemented with 2 mM glutamine. Basal ECAR was recorded, followed by the sequential injection of 10 mM glucose, 1 μM oligomycin, and 50 mM 2-DG. The obtained data were then normalized to the total protein content per well.

#### Measurements of lactate concentrations

Lactate levels in the cell culture medium of VSMCs were measured using the colorimetric lactate assay kit (Jiancheng Bioengineering Institute, Nanjing, China) according to the manufacturer’s instructions. The data were then normalized to the total protein concentration, which was determined using the BCA Protein Assay Kit.

#### Molecular docking

The crystal structure of PFKFB3 was retrieved from the Protein Data Bank (PDB) with PDB ID: 6ETJ. The 3D structure of Liriodendrin (Lidd) was obtained from PubChem (PubChem CID: 21603207), and energy minimization of the ligand was performed using Chem3D under the MMFF94 force field. The Molecular Operating Environment (MOE) software (2020 version) was used to analyze the interaction of PFKFB3 with Lidd. PFKFB3 was set as receptor and Lidd as the ligand. When PFKFB3-Lidd docking finished, the lowest energy postures with the strongest binding were selected in Database Viewer to search for the most likely binding regions and binding sites of protein-ligand. Fingerprint analysis in MOE was used to further evaluate the docking results.

#### Drug affinity responsive target stability (DARTS)

Cell lysates were prepared by lysing untreated hPASMCs on ice for 30 minutes using a mammalian protein extraction reagent (Thermo Scientific). Equal amounts of protein (500 μg per aliquot, 5 μg/μL) were incubated with or without the indicated concentrations of Liriodendrin (Lidd) in 1 × TNC buffer (50 mmol/L Tris, 50 mmol/L NaCl, 10 mmol/L CaCl2, pH 7.4) at room temperature for 2 hours. Proteinase K from Tritirachium album (Sigma-Aldrich) was then simultaneously added to all samples at a 1:1000 (w/w) ratio and incubated at 25°C for 15 minutes. Following digestion, loading buffer was added to each sample, and the lysates were boiled for 10 minutes. The resulting samples were subjected to western blotting analysis.

#### Western blot analysis

Proteins were extracted from lung homogenates, or cells using RIPA lysis buffer supplemented with a phosphatase inhibitor cocktail. The protein concentration was determined using the BCA Protein Assay Kit (Vazyme, Nanjing, USA). Equal amounts of protein (20-50 μg) were separated by sodium dodecyl sulphate-polyacrylamide gel electrophoresis and transferred onto a polyvinylidene difluoride (PVDF) membrane. The membranes were blocked for 1 hour at room temperature with 5% fat-free milk in Tris-buffered saline containing 0.1% Tween 20 (TBST). After blocking, the membranes were incubated overnight at 4°C with primary antibodies. Following three washes with TBST, the membranes were incubated for 1 hour at room temperature with HRP-conjugated secondary antibodies (anti-rabbit or anti-mouse IgG, according to the origin of the primary antibodies). Protein bands were visualized using an enhanced chemiluminescence reagent (Thermo Fisher Scientific, Waltham, MA, USA), and the band intensities were quantified using Fiji software. The protein expression levels were normalized to the loading control for comparative analysis.

#### Cell transfection

Cells were plated in 6-well plates to achieve 50-70% confluency. siRNA or plasmid and jetPRIME transfection reagent were mixed and incubated at room temperature for 10 minutes to form transfection complexes. The complexes were then added to the cells, and incubated for 12 hours. Afterward, the transfection mixture was removed, and the cells were cultured under standard conditions. Knockdown and overexpression efficiencies were evaluated 48 hours post-transfection using RT-qPCR or Western blotting.

#### Immunoprecipitation

After centrifugation and washing with PBS, the cell pellets were resuspended and lysed using immunoprecipitation lysis buffer (Beyotime, Shanghai, China) following the manufacturer’s protocol. One milligram of total cell lysates was incubated overnight at 4°C with gentle shaking using 10 μL of anti-FLAG magnetic beads (Selleck). The beads were then washed three times with immunoprecipitation washing buffer (150 mM Tris-HCl, 400 mM NaCl, 0.8% Triton X-100, pH 7.4), boiled in 2× loading buffer, and subjected to Western blot analysis.

#### Chromatin immunoprecipitation (ChIP)

ChIP was performed using the SimpleChIP Enzymatic Chromatin IP Kit (CST, USA) according to the manufacturer’s instructions. Briefly, cells were crosslinked at room temperature by incubating with 2 mM disuccinimidyl glutarate for 45 minutes, followed by 1% formaldehyde for 10 minutes. Chromatin was then isolated and sonicated to achieve fragment sizes of 200-500 bp. The samples were incubated overnight at 4°C with rotation using antibodies specific to H3K18la or normal rabbit IgG as a negative control. The immunoprecipitated DNA fragments were subsequently captured, eluted, and crosslinking was reversed. The purified DNA was quantified using real-time qPCR with specific primers targeting the promoters of the indicated genes (primer sequences are provided in **Table S4**).

#### Statistical analysis

All statistical analyses were performed using GraphPad Prism (version 8.0; GraphPad Software, San Diego, CA, USA). Data are presented as the mean ± SEM from independent biological replicates. The normality of data distribution was assessed using the Shapiro-Wilk test. For comparisons between two groups, an unpaired two-tailed Student’s t-test was used for normally distributed data, while the Mann–Whitney U test was applied for non-normally distributed data. For multiple-group comparisons, one-way or two-way analysis of variance (ANOVA) followed by Tukey’s post hoc test was used for normally distributed data, whereas Welch’s ANOVA followed by Dunnett’s T3 test was used for non-normally distributed data. The specific statistical tests used for each analysis are detailed in the figure legends. A *p*-value < 0.05 was considered statistically significant.

## Acknowledgements

This work was supported by the Noncommunicable Chronic Diseases National Science and Technology Major Project (grant: 2024ZD0528700), National Natural Science Foundation of China (grants: 82422002) and Shanghai Science and Technology Commission of China (grant: 20410761000).

## Author contributions

D.K., S.L. and Z.L. conceptualized the study; Q.Z., Z.D., L.Y., and J.S. developed the methodology; Q.Z., Z.D., Q.L., Z.S., Y.L., X.H. and J.Z. performed data analysis; Q.Z., Z.D. and Q.L. created the visualization; D.K. and Z.L. acquired the funding; D.K. S.L. and Z.L. administered the project; D.K. and Z.L. supervised the research; D.K. Q.Z. and Z.D. wrote the original draft; D.K., Z.D., Z.L. and S.L. reviewed and edited the manuscript.

## Abbreviations

α-SMA: a-Smooth muscle actin
CCL2: C-C Motif Chemokine Ligand 2
CCND1: D1cyclin D1
CUT&Tag: Cleavage Under Targets and Tagmentation
CXCL1 C-X-C: Motif Chemokine Ligand 1
EdU: 5-ethynyl-2′-deoxyuridine
FZR1: Fizzy-Related Protein Homolog
H3K18la: histone H3 lysine 18 lactylation
Lidd: Liriodenrin
MCT: pmonocrotaline
PASMCs: arterialpulmonary arterial smooth muscle cells
PAH: pulmonary arterial hypertension
PCNA: proliferating cell nuclear antigen
PDGF-BB: platelet-derived growth factor-BB
PFKFB3: 6-phosphofructo-2-kinase/fructose-2,6-bisphosphatase 3
PH: pulmonary hypertension
RV/(LV+S): ratio of right ventricular weight to the left ventricle plus septal weight
RVSP right: ventricular systolic pressure
SuHx: Sugen 5416 plus hypoxia
Ub: Ubiquitin

---

**Figure S1.**
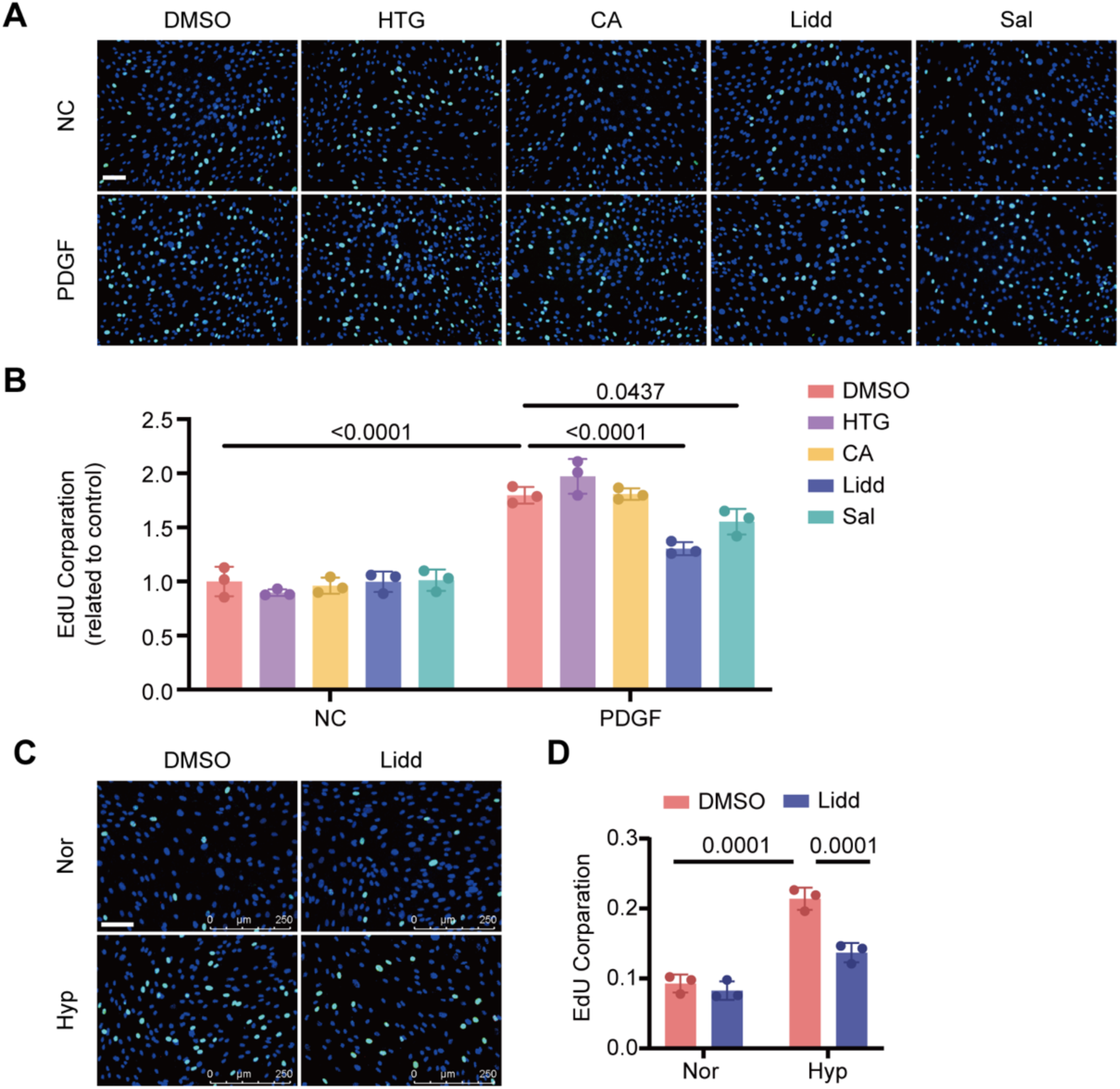
Effect of four active ingredients on pulmonary artery smooth muscle cell (PASMC) proliferation. **A-B**) Representative EdU assay (**A**) and quantification of EdU incorporation (**B**) in hPASMCs treated with PDGF-BB (20 ng/mL) or vehicle in the presence of 100 μM hydroxytyrosol-1-glucopyranoside (HTG), chlorogenic acid (CA), liriodendrin (Lidd), or salidroside (Sal) for 48 h. Scale bar, 100 μm (n = 3). **C-D**) Representative EdU assay (**C**) and quantification of EdU incorporation (**D**) in hPASMCs cultured under normoxia (Nor) or hypoxia (Hyp) and treated with DMSO or Lidd. Scale bar, 100 μm (n = 3). Data are presented as mean ± SEM. Statistical analyses were performed using two-way ANOVA followed by Tukey’s multiple-comparisons test. NC, negative control; PDGF, platelet-derived growth factor-BB; EdU, 5-ethynyl-2ʹ-deoxyuridine; Nor, normoxia; Hyp, hypoxia.

**Figure S2.**
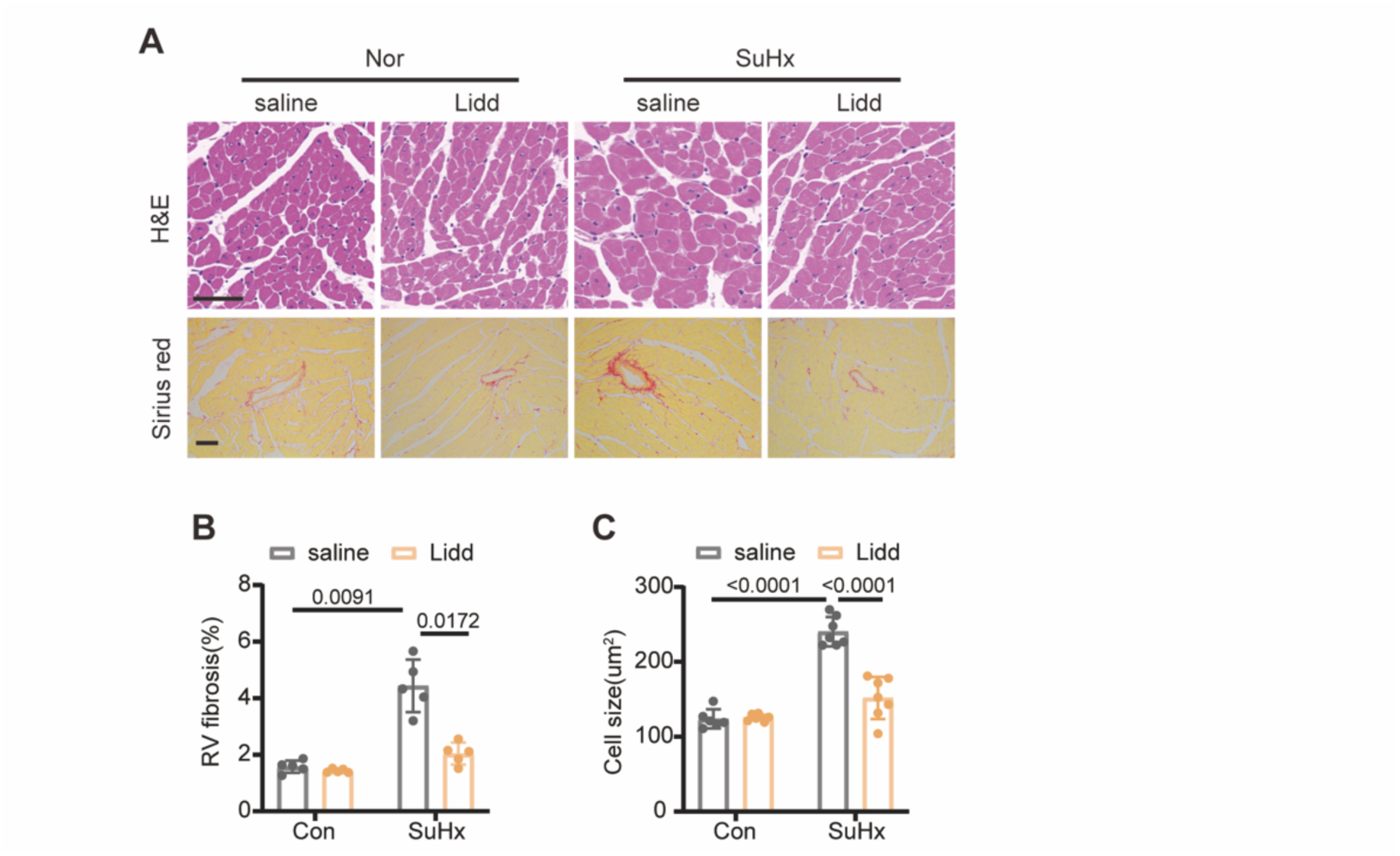
Lidd attenuates right ventricular remodeling in SuHx-induced pulmonary hypertension. **A**) Representative images of hematoxylin and eosin (H&E) staining (**top**) and Sirius red staining (**bottom**) in right ventricular sections from Nor- or SuHx-exposed mice treated with saline or Lidd. Scale bars, 50 μm. **B-C**) Quantification of right ventricular fibrosis (**B**) and cardiomyocyte cross-sectional area (**C**) in **A** (n ≥ 5). Data are presented as mean ± SEM. Statistical analyses were performed using two-way ANOVA followed by Tukey’s multiple-comparisons test. Nor, normoxia; SuHx, SU5416/hypoxia.

**Figure S3.**
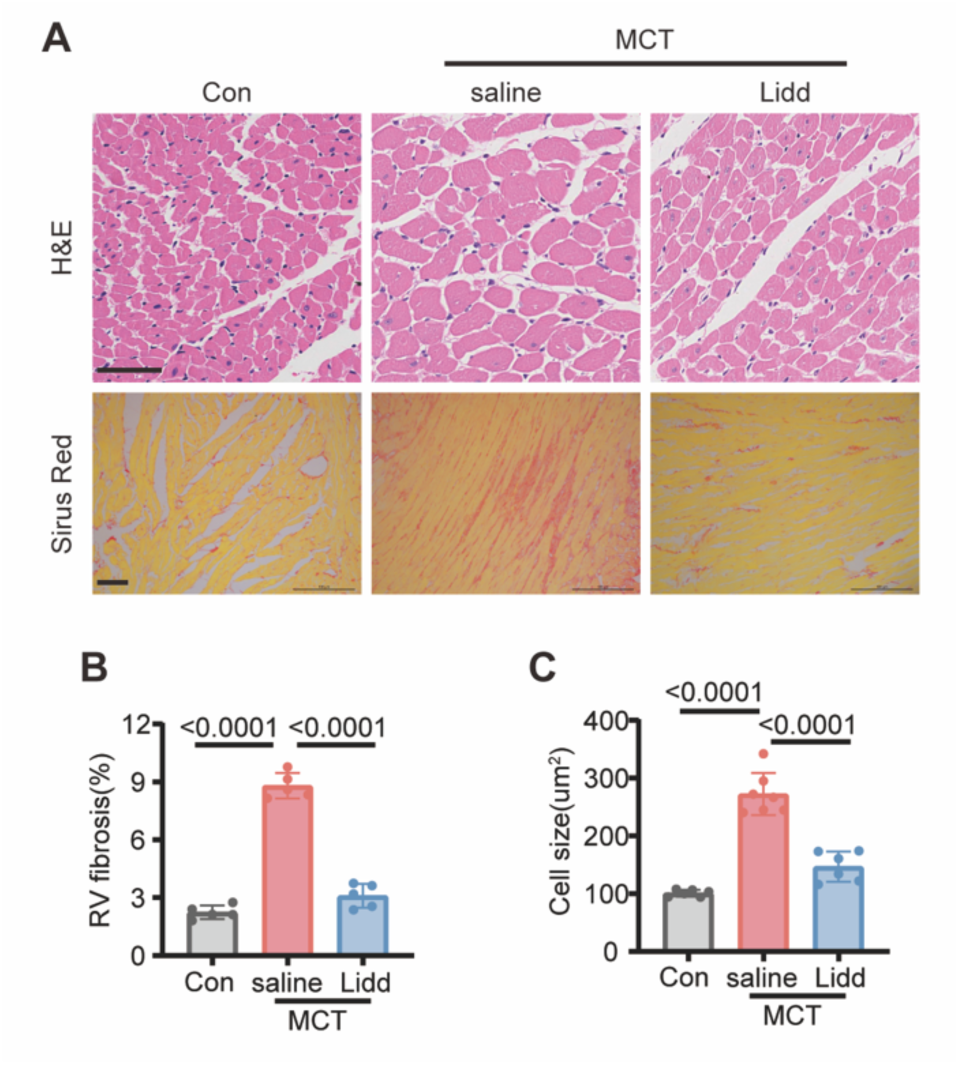
Lidd attenuates right ventricular remodeling in MCT-induced pulmonary hypertension. **A)** Representative images of hematoxylin and eosin (H&E) staining (**top**) and Sirius red staining (**bottom**) in right ventricular sections from control or MCT-treated rats with or without Lidd treatment. Scale bars, 50 μm. **B-C**) Quantification of right ventricular fibrosis (**B**) and cardiomyocyte cross-sectional area (**C**) in **A** (n ≥ 5). Data are presented as mean ± SEM. Statistical analyses were performed using one-way ANOVA followed by Tukey’s multiple-comparisons test. Con, vehicle control; MCT, monocrotaline.

**Figure S4.**
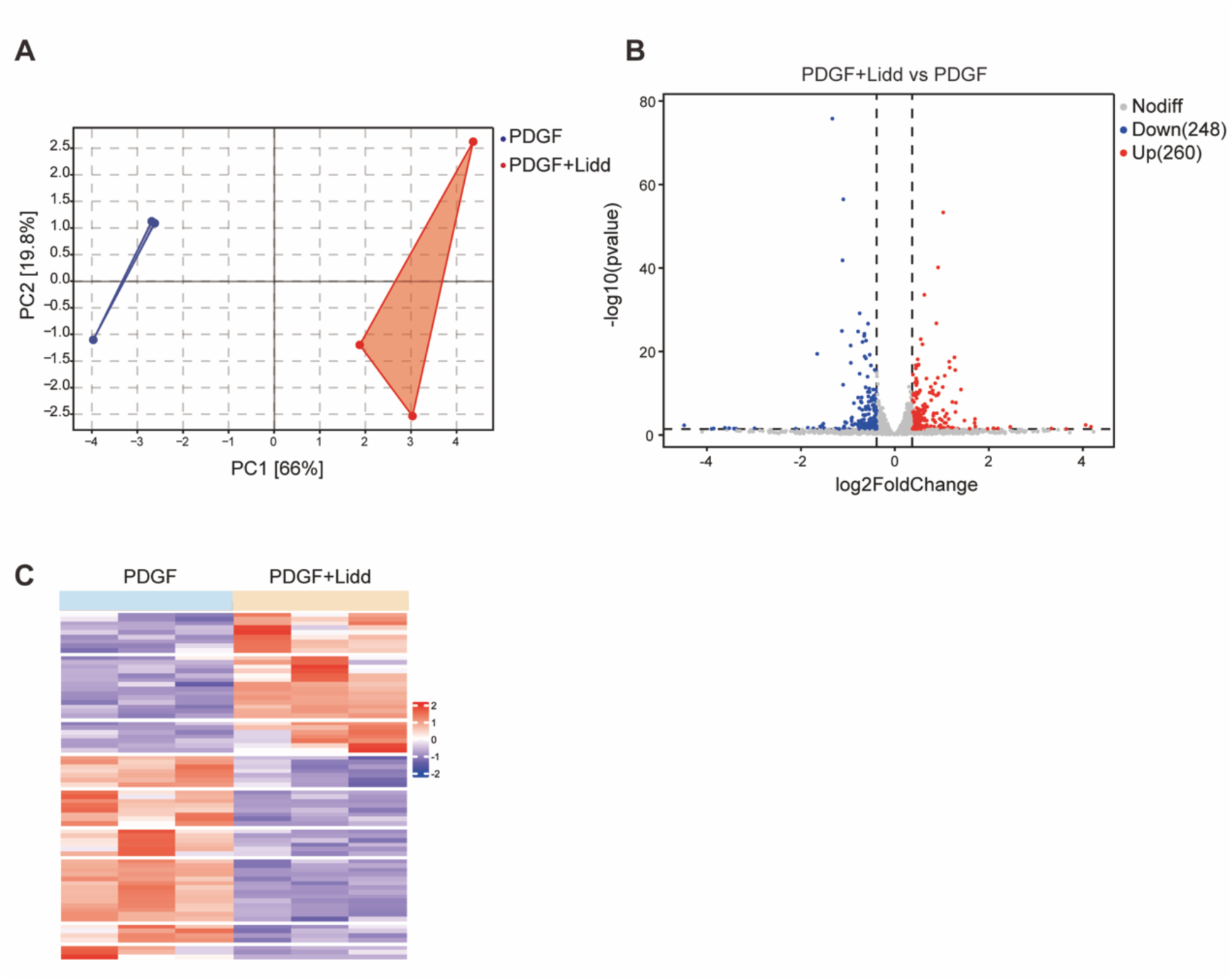
Characteristics of RNA-seq data from PDGF-BB-stimulated human PASMCs (hPASMCs) with or without Lidd treatment. hPASMCs were treated with PDGF-BB (20 ng/mL) in the presence of DMSO or Lidd (100 μM) for 24 h. A) Principal component analysis (PCA) of RNA-seq data from PDGF-BB-stimulated hPASMCs treated with DMSO or Lidd (n = 3). B) Volcano plot of differentially expressed genes (DEGs) in PDGF-BB-stimulated hPASMCs treated with Lidd or DMSO. Differentially expressed genes were defined as genes with an absolute fold change ≥ 0.58 and an adjusted P value (Q value) ≤ 0.001. C) Heatmap of differentially expressed genes in PDGF-BB-stimulated hPASMCs treated with DMSO or Lidd for 24 h (n = 3).

**Figure S5.**
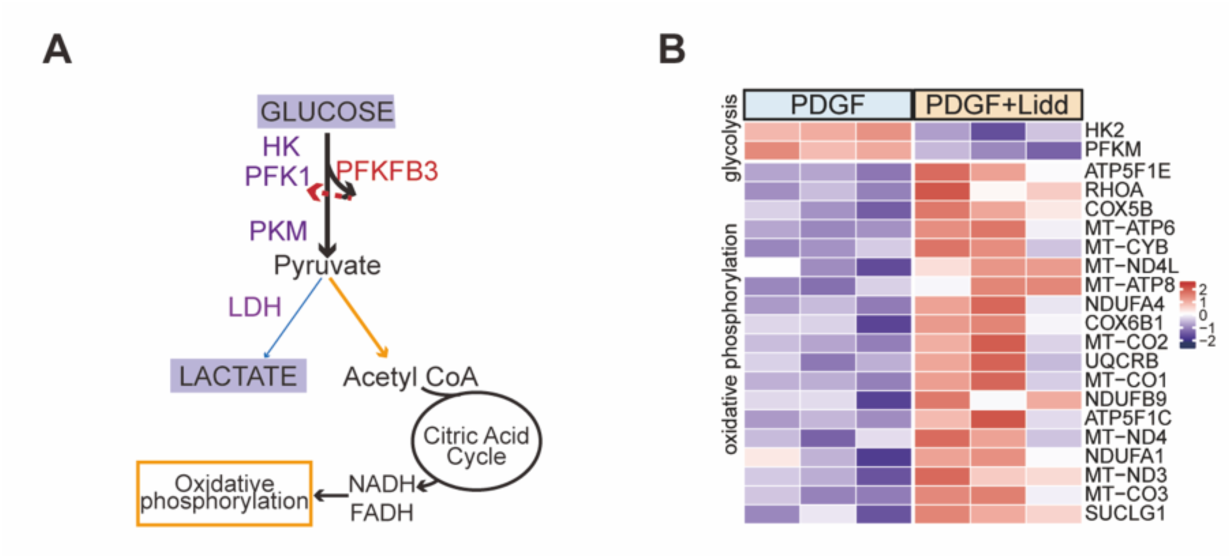
Lidd suppresses PDGF-BB-induced glucose metabolism reprogramming in hPASMCs. **A)** Schematic representation of key enzymes and products in glycolysis and oxidative phosphorylation (OXPHOS), the two primary ATP-producing pathways. **B)** Heatmap of differentially expressed genes associated with glycolysis and oxidative phosphorylation (OXPHOS) identified by RNA sequencing of PDGF-BB-stimulated hPASMCs treated with DMSO or Lidd for 24 h (n = 3).

**Figure S6.**
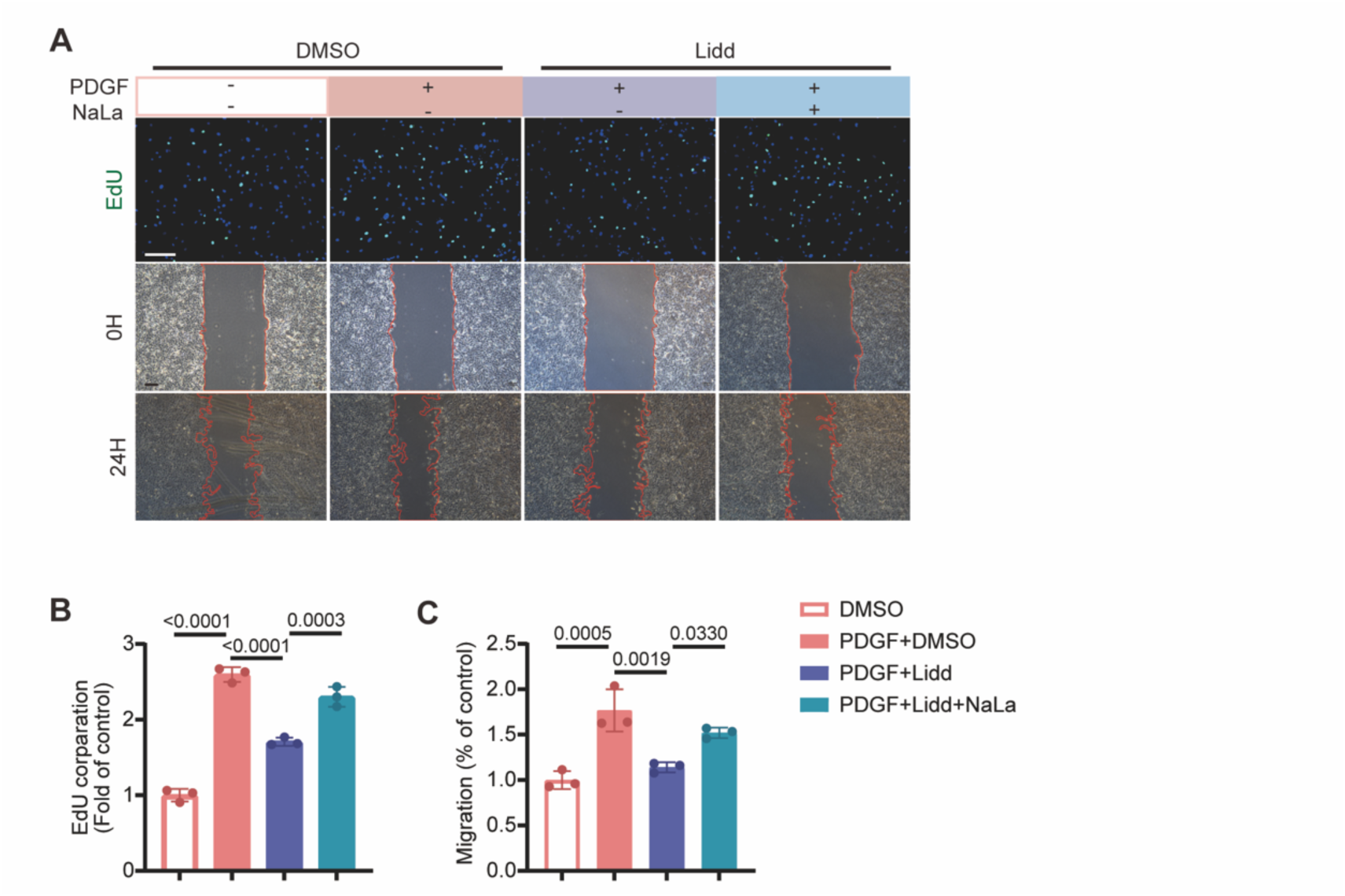
Supplementation with sodium lactate restores Lidd-induced suppression of proliferation and migration in PDGF-BB-stimulated hPASMCs. **A-C)** Representative EdU assay (**top**) after 48 h of treatment and wound healing assay (**bottom**) after 24 h of treatment (**A**), and quantification of EdU incorporation (**B**) and wound closure (**C**) in hPASMCs treated with PDGF-BB or vehicle in the presence of DMSO or Lidd. Sodium lactate (NaLa, 5 mM) was added where indicated. Scale bars, 200 μm (n = 3). Data are presented as mean ± SEM. Statistical analyses were performed using one-way ANOVA followed by Tukey’s multiple-comparisons test. NaLa, sodium lactate; EdU, 5-ethynyl-2′-deoxyuridine.

**Figure S7.**
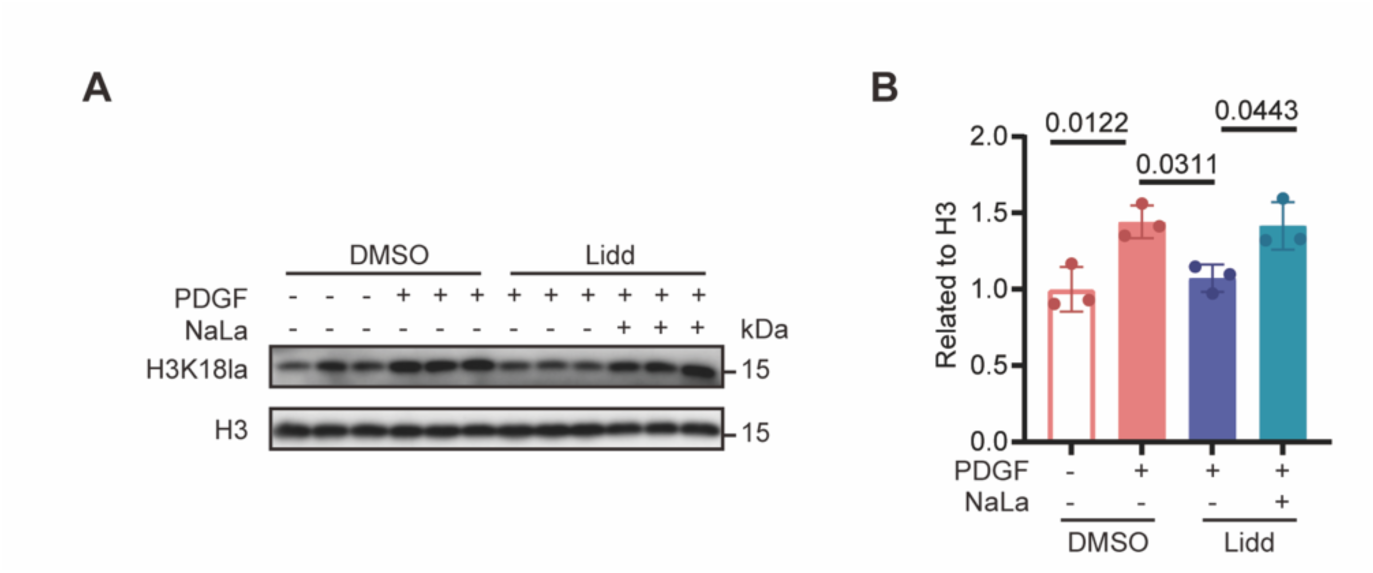
Sodium lactate (NaLa) reverses Lidd-mediated inhibition of histone H3 lysine 18 lactylation (H3K18la). **A-B)** Representative immunoblotting (**A**) and quantification (**B**) of H3K18la levels in hPASMCs treated with PDGF-BB or vehicle in the presence of DMSO or Lidd. Sodium lactate (NaLa, 5 mM) was added where indicated Sodium lactate (n = 3). Data are presented as mean ± SEM. Statistical analyses were performed using one-way ANOVA followed by Tukey’s multiple-comparisons test. NaLa, sodium lactate; PDGF, platelet-derived growth factor-BB; H3K18la, histone H3 lysine 18 lactylation; H3, histone H3.

**Figure S8.**
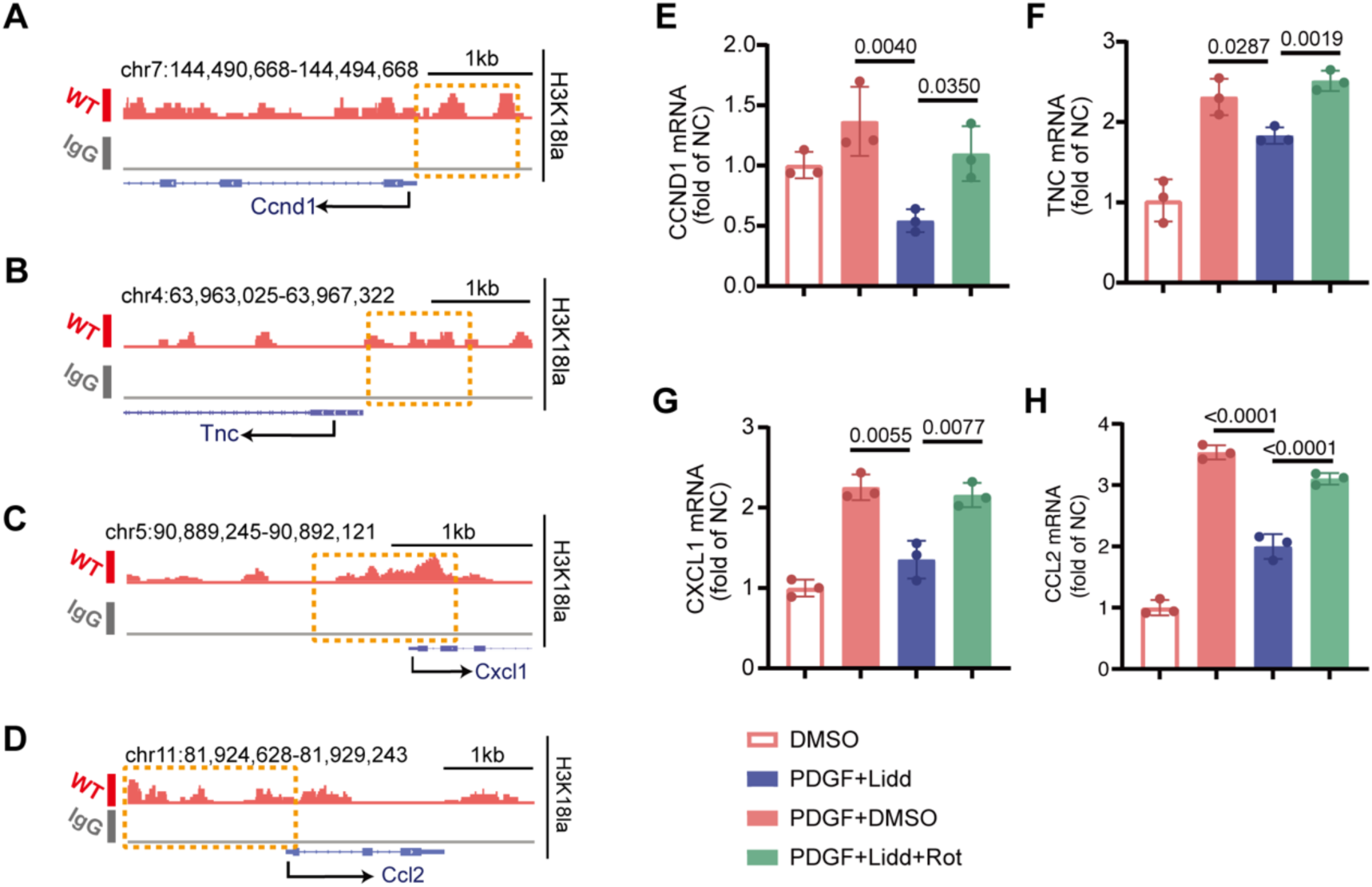
H3K18la enrichment at the Ccnd1, Tnc, Cxcl1 and Ccl2 loci and regulation of their expression by Lidd. **A-D)** Representative Integrative Genomics Viewer (IGV) tracks of publicly available H3K18la CUT&Tag-seq data showing H3K18la enrichment at the Ccnd1 (**A**), Tnc (**B**), Cxcl1(**C**) and Ccl2 (**D**) loci in synthetic VSMCs. **E-H)** qRT-PCR analysis of CCND1 (**E**), TNC (**F**), CXCL1(**G**) and CCL2 (**H**) mRNA expression in hPASMCs treated with PDGF-BB or vehicle in the presence of DMSO or Lidd. Rotenone (Rot, 5 nM) was added where indicated (n = 3). Data are presented as mean ± SEM. Statistical analyses were performed using two-way ANOVA followed by Tukey’s multiple-comparisons test. IGV, Integrative Genomics Viewer; Rot, rotenone; H3K18la, histone H3 lysine 18 lactylation.

**Figure S9.**
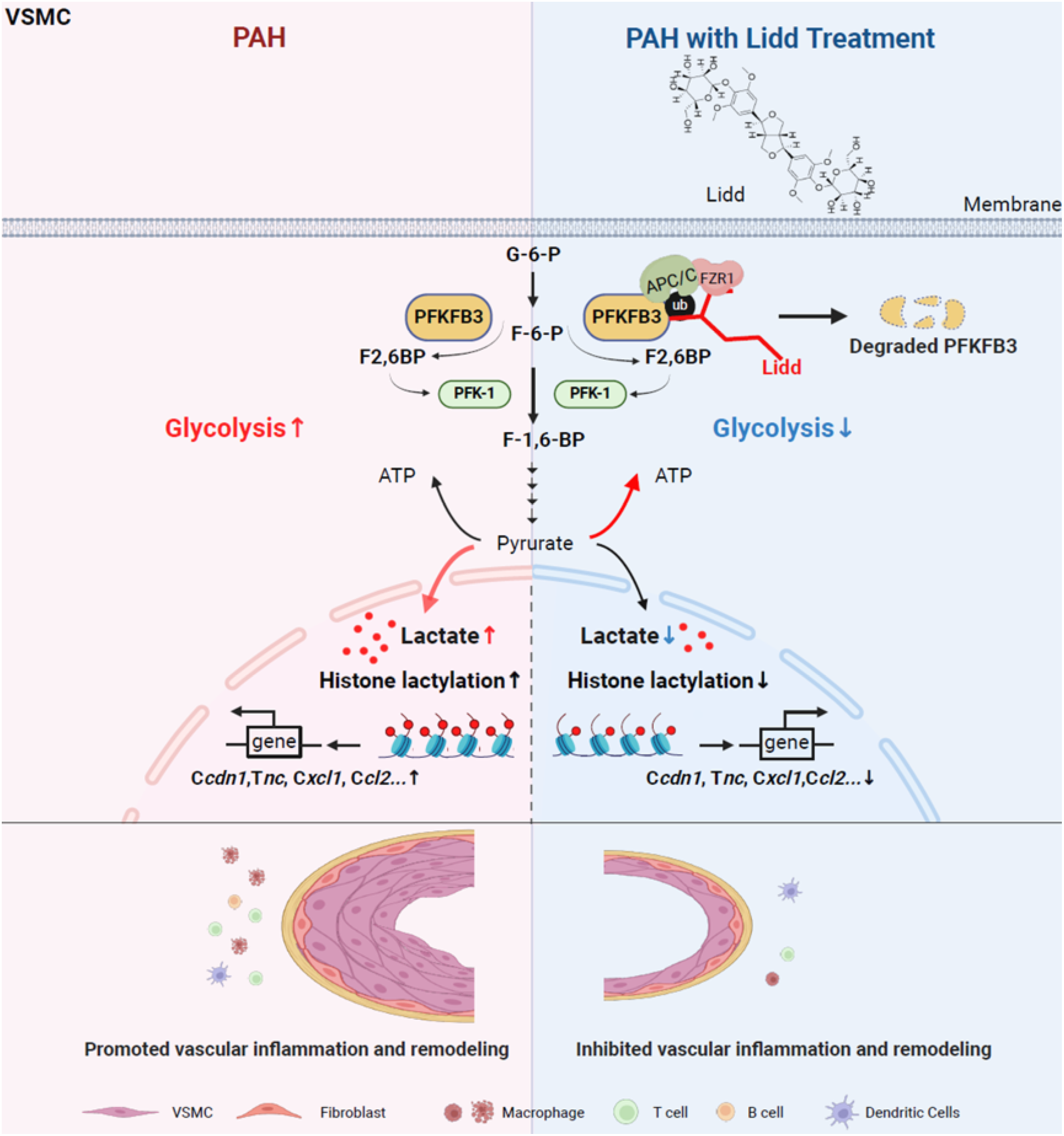
Proposed mechanism by which Lidd attenuates pulmonary vascular remodeling in pulmonary arterial hypertension. Schematic illustrating the proposed mechanism of action of Lidd in pulmonary arterial hypertension (PAH). In PAH, increased PFKFB3-driven glycolysis enhances lactate production and histone H3 lysine 18 lactylation (H3K18la), leading to transcriptional activation of genes associated with pulmonary artery smooth muscle cell phenotypic transformation, inflammation, and vascular remodeling. Lidd promotes FZR1-mediated ubiquitination and proteasomal degradation of PFKFB3, thereby suppressing glycolysis, reducing lactate production and H3K18la, and attenuating pulmonary vascular inflammation and remodeling.

**Figure S10.**
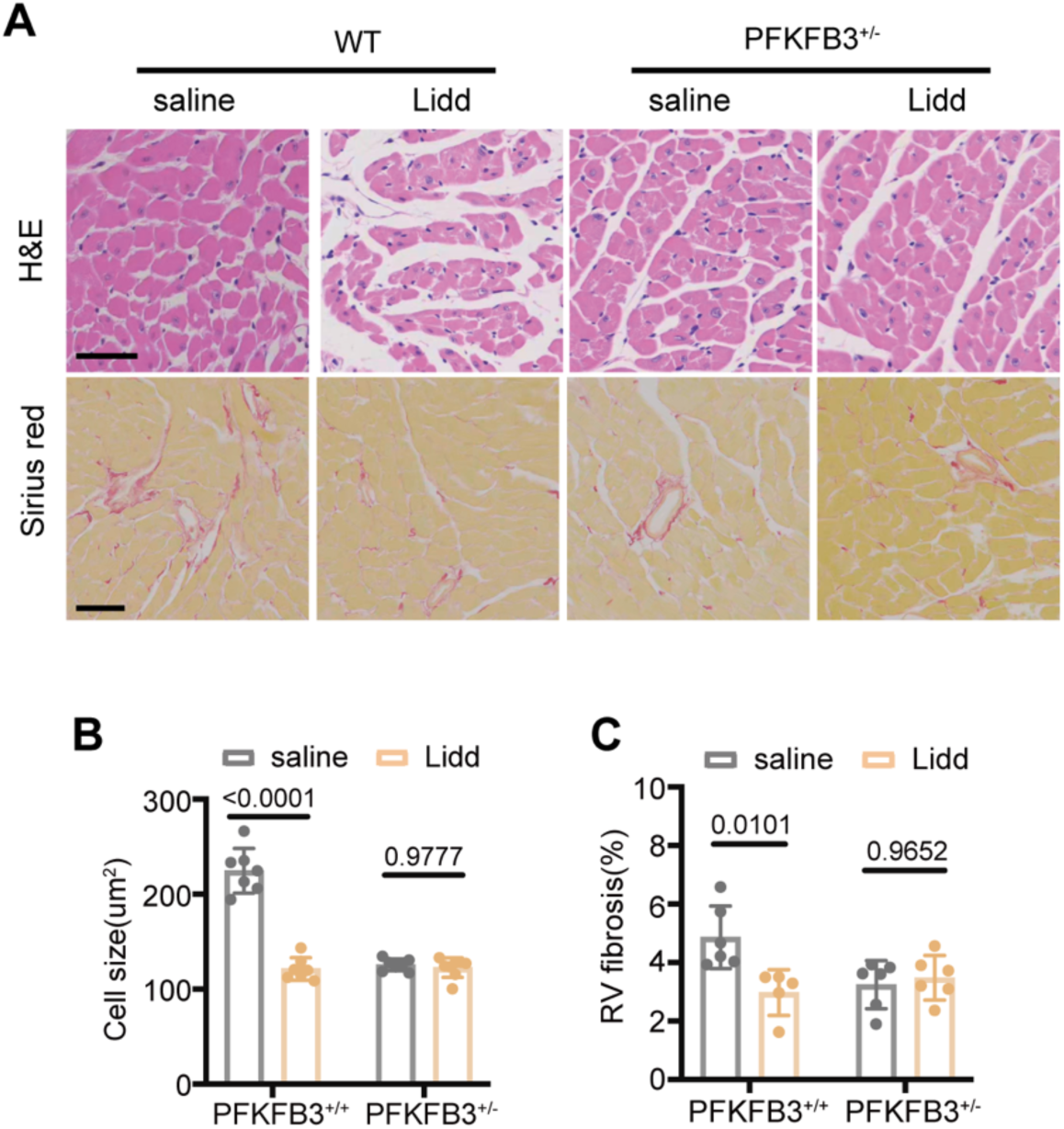
PFKFB3 deficiency abolishes the protective effects of Lidd on right ventricular remodeling in SuHx-induced pulmonary hypertension. **A)** Representative images of hematoxylin and eosin (H&E) staining (**top**) and Sirius red staining (**bottom**) in right ventricular sections from SuHx-exposed PFKFB3^+/+^ and PFKFB3^+/-^ mice treated with saline or Lidd. Scale bars, 50 μm. **B–C)** Quantification of cardiomyocyte cross-sectional area (**B**) and right ventricular fibrosis (**C**) in **A** (n ≥ 5). Data are presented as mean ± SEM. Statistical analyses were performed using two-way ANOVA followed by Tukey’s multiple-comparisons test. SuHx, SU5416/hypoxia; RV, right ventricle.

